# LAmbDA: Label Ambiguous Domain Adaption Dataset Integration Reduces Batch Effects and Improves Subtype Detection

**DOI:** 10.1101/522474

**Authors:** Travis S Johnson, Zhi Huang, Christina Y Yu, Tongxin Wang, Yi Wu, Yatong Han, Kun Hung, Jie Zhang

## Abstract

**Motivation:** Rapid advances in single cell RNA sequencing have produced more granular subtypes of cells in multiple tissues from different species. There exists a need to develop rigorous methods that can i) model multiple datasets with ambiguous labels across species and studies and ii) remove systematic biases across datasets and species.

**Results:** We developed a species- and dataset-independent transfer learning framework (LAmbDA) to train models on multiple datasets and applied our framework on scRNA-seq experiments. These models mapped corresponding cell types between datasets with inconsistent labels while simultaneously reducing batch effects. We achieved high accuracy in labeling cellular subtypes (weighted accuracy pancreas: 91%, brain: 78%) using LAmbDA Random Forest. LAmbDA Feedforward 1 Layer Neural Network achieved higher weighted accuracy in labeling cellular subtypes than CaSTLe or MetaNeighbor in brain (48%, 32%, 20% respectively). Furthermore, LAmbDA Feedforward 1 Layer Neural Network was the only method to correctly predict ambiguous cellular subtype labels in both pancreas and brain compared to CaSTLe and MetaNeighbor. LAmbDA is model- and dataset-independent and generalizable to diverse data types representing an advance in biocomputing.

**Availability:** github.com/tsteelejohnson91/LAmbDA

## 1 Introduction

Amidst trillions of cells and hundreds of distinct cell types in the human body, understanding tissue heterogeneity and the resulting phenotypic consequences is a mammoth task with far-reaching impact. For example, the brain consists of diverse co-localized neural, glial, immune, and vascular cell types that work in concert to form complex nervous tissues. Complex tissues and their constituent cell types have already been studied at the tissue level of granularity (Dorrell, et al., 2008; Dorrell, et al., 2011; Erlandsen, et al., 1976; Gomori, 1939; Zhang, et al., 2014). Fundamentally, these tissues are composed of intricate populations of cells; researchers are now turning to the single cell level to discern new cellular subtypes (Baron, et al., 2016; Darmanis, et al., 2015), which are often spatially indistinct in their tissue of origin (Kumar, et al., 1999). For these reasons, there is a critical need to differentiate cells from complex tissues during sequencing.

The rapid advance of single cell RNA sequencing (scRNA-seq) enables researchers to study cell differentiation and tissue heterogeneity in various, tissues, diseases and physiological states. Studies have analyzed scRNA-seq data from different species, such as mouse (Chen, et al., 2017; Li, et al., 2016; Zeisel, et al., 2015) and human (Darmanis, et al., 2015; Lake, et al., 2016). Tissue studies have conducted mouse-human comparisons (Baron, et al., 2016) and normal-diabetes comparisons (Segerstolpe, et al., 2016). Some studies have directly compared human and mouse cell types from the same brain region (Johnson, et al., 2016; La Manno, et al., 2016). These studies are especially important if data from mouse tissues can be used to identify or fill in the missing human tissues of counterpart cell types into “*in silico* chimeric” datasets. These integrative datasets could prove especially useful when human data is scarce or technically infeasible to generate. However, the increased number of scRNA-seq experiments has produced unforeseen challenges.

One such challenge arises in that each scRNA-seq dataset generates its own subtype labels, which are often identified based on unsupervised approaches, such as clustering, and carry intrinsic systemic biases (i.e. batch effects). These labels are often not consistent enough to be directly used across datasets/studies/species without first identifying their correspondence to each other. There have been efforts to i) identify the correspondence of subtypes across datasets using gene set correlations (Crow, et al., 2018), ii) to combine datasets for integrative clustering (Butler, et al., 2018), and iii) predict labels in one dataset with another (Lieberman, et al., 2018). These represent three of the major tasks in combining scRNA-seq datasets for analysis. The second task is significant in that it can remove batch effects when clustering single cells from multiple experiments (Butler, et al., 2018; Lin, et al., 2018; Risso, et al., 2018; Zappia, et al., 2018). However, these methods often require labels to have a precise match between datasets and none of these methods address all three tasks simultaneously. The third methodology leverages transfer learning, a subset of machine learning, but cannot simultaneously train on more than two datasets.

In transfer learning, neural networks (NNs) can be trained more efficiently and effectively on a target task when first trained on source examples (Pratt, 1993). Training on multiple datasets drawn from different distributions can reduce the amount of sample selection bias, a potential cause of batch effects, in the resulting model (Huang, et al., 2006). Furthermore, unknown labels can be derived through domain adaptive training, resulting in a target task with labels (Ganin, et al., 2016). In computer vision, there have been multiple studies aiming at training convolutional NNs with label ambiguity (Cour, et al., 2011; Geng 2017; Hullermeier and Beringer, 2005; Jie and Orabona, 2010).

Fortunately, recent developments in deep learning have allowed NNs to accomplish classification and identification tasks in scRNA-seq. For example, (Chu, et al., 2016) leveraged the large amount of scRNA-seq data to train NN classifiers and identified the tissues of origin in circulating cells. However, these NN models, while important for feature reduction and identifying tissue of origin, were not optimally trained to be accurate across species in a single tissue type (Lin, et al., 2017) and did not carry out dataset integration with other tissues despite the data rich environment of single cell transcriptomics (Andrews and Hemberg, 2018). To take advantage of single-cell data from different sources and species, effective machine learning algorithms are needed for across-species cell type mapping and gene feature reduction.

In this paper, we present a novel integrative transfer learning framework called LAmbDA (<u>L</u>abel <u>Amb</u>iguous <u>D</u>omain <u>A</u>daption), which reduces inter-dataset distances and learns the label for ambiguously labeled cells. We tested multiple machine learning algorithms including logistic regression (LR), Feedforward 1 Layer NN (FF1), Feedforward 3 Layer NN (FF3), Recurrent Neural Network (RNN1), and Random Forest (RF) to optimize LAmbDA, and applied it to both human pancreas and human/mouse brain scRNA-seq datasets for subtype identification and matching. Subtypes of cells shared across datasets are considered replicable and robust (Crow, et al., 2018). We refer to these robust classes of cellular subtypes as “conserved” since they are consistent regardless of dataset, species, and condition. These biologically relevant conserved subtypes were discovered by LAmbDA.

To summarize, we demonstrate that LAmbDA-based models are capable of simultaneously matching unstandardized labels with varying degrees of overlap, combining disparate datasets from different species/platforms using training and testing set, and predicting conserved subtypes of cells learned during training with high accuracy. LAmbDA can serve as the framework to accommodate other models beyond these biological applications to suit a variety of data types and analyses.

## 2 Methods

### 2.1 Datasets

Six scRNA-seq datasets were used to test LAmbDA in two different tissue types consisting of three pancreatic and three brain scRNA-seq datasets. We intentionally chose a heterogeneous mix of datasets to study the robustness of our method.

The pancreatic datasets included (Fig. 1A): one human dataset with 15 cell types (Seg, 1980 cells) (Segerstolpe, et al., 2016), one human dataset with 10 cell types (Mur, 2126 cells) (Muraro, et al., 2016), and one human dataset with 14 cell types (Bar, 8569 cells) (Baron, et al., 2016). The brain datasets included (Fig. 1B): one human dataset with only neurons and 16 subtype level labels (HumN, 3086 cells) (Lake, et al., 2016), one human dataset with neurons and glia and six major cell type level labels (HumNG, 285 cells) (Darmanis, et al., 2015), and one mouse dataset with neurons and glia and 48 subtype level labels (MusNG, 3005 cells) (Zeisel, et al., 2015).

**Fig. 1.**
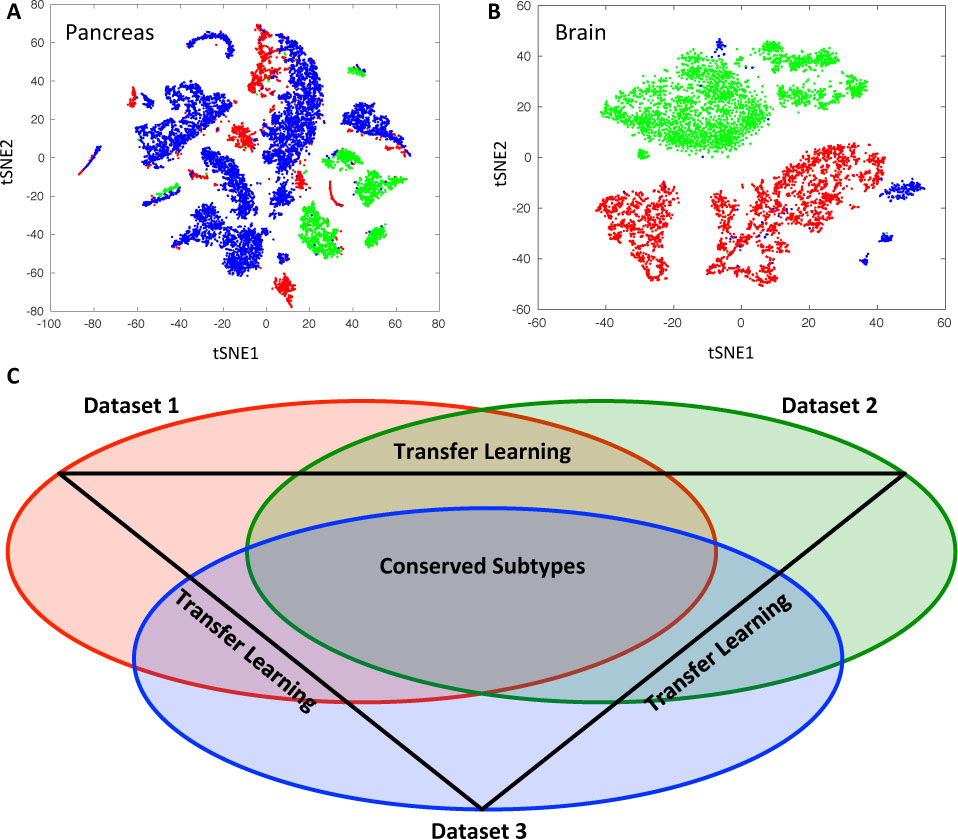
t-SNE plot of scRNAs-seq data after feature selection step. A) Pancreatic datasets. Colors indicate different datasets: Seg (red), Mur (green), Bar (red). B) Brain datasets: MusNG (red), HumN (green), HumNG (blue). C) A scheme of conserved subtype identification using transfer learning approach (a three-dataset example).

### 2.2 General Framework

#### Dataset Integration

We illustrate the LAmbDA framework using an example with three different datasets. In our annotation, bold uppercase denotes matrix (**X**), bold lowercase denotes vector (**x**), lowercase letter denotes numeric value (x), and uppercase denotes a set (e.g. gene set or sample set, **X**). Given three scRNA-seq expression matrices (***X*_1_**, ***X*_2_**, ***X*_3_**) with *n*_1_, *n*_2_, *n*_3_ cells (samples) and *T*_l_, *T*_2_, *T*_3_ transcripts (feature) sets, the number of transcripts are first reduced to the intersection of all three datasets (*T*). The subtype labels of each cell across all three datasets are denoted by ***Y*_1_, *Y*_2_, *Y*_3_** each containing *l*_1_, *l*_2_, *l*_3_ labels, respectively, shown below:

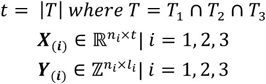

The labels are one-hot encoded such that each row of ***Y***_(***i***)_ contains a single value of one indicating the label of the specific cell. Each row will have a single value of one in the column corresponding to that subtype label. To pool all of the datasets together for a single model, we combine the expression matrix (***X***) and label matrix (***Y***) described below:

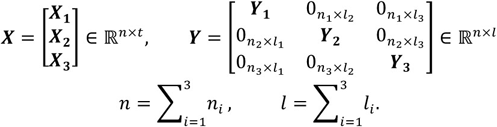

Using this encoding, it would be straightforward to train a logistic regression, random forest, or NN model (*f*(***X***)) on the data using one of the multiple optimization algorithms to minimize the following objective function:

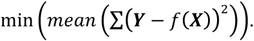

However, all the labels (*L*) in the three datasets are not identical nor mutually exclusive. For example, in the brain study, all interneuron subtypes in dataset 2 could potentially match any of the interneuron subtypes in dataset 1. This label overlap between datasets means a subset of the more refined conserved subtypes (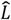) exists in *L* such that all sub-types in 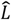 can be assigned to a subtype in *L* (Fig. 1C). A new and more refined label matrix (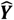) is generated using 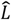:

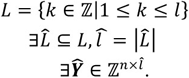

As a result, we propose that it is possible to train a model 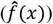 on the more refined subtypes (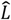 and 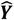) using an optimization algorithm on the following optimization problem:

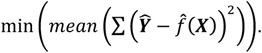

The above optimization problem is solved using the two following algorithms. Algorithm 1 corresponds to the more general version of LAmbDA used for LR and RF. Algorithm 2 corresponds to the NN implementation that actively removes batch effects in the hidden layer.

#### Algorithms

To train the LAmbDA models, we used the Adam Optimizer (Kingma and Ba, 2014) with step size of 0.01 and random mini-batches of size ***p_batch_*** (a percentage of 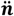, see **Eq S3**) that were changed every 50 iterations to prevent overfitting of unambiguous labels. We ran each model for 2000 iterations except for the RF model, which was run for 100 iterations. The code was written for GPU-enabled TensorFlow Python3 package. The input matrices (***X*, *Y***) were preprocessed into 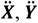 and the possible inter-dataset label mappings were preprocessed into an adjacency matrix (***G***) before running the algorithms. For details on the preprocessing and individual equations used, please see **Supplementary Material Sec. 2.1**.

##### Algorithm 1 Label Ambiguous Domain Adaption (LAmbDA)

**Input:** preprocessed expression matrix 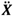, preprocessed labels 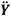, and label mask *G***, Eq S1-3**

**Output:** a trained classifier 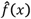 with mapped ambiguous labels and batch effects removed Random initialization

**1. Train on unambiguous labels** Using the subset of samples that have only one possible label For the first half of total iterations:

**i. Forward propagate** predicted labels
**ii. Back propagate** gradient from label error (i.e. update model)

**2. Train on ambiguous labels** Using all samples regardless of number of possible labels For the second half of total iterations:

**i. Forward propagate** predicted labels (i.e. calculate 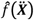, **Eq S13,17**)
**ii. Assign labels** to ambiguously labeled cells (i.e. calculate 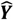, **Eq S8**)
**iii. Calculate label error** using 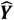 and 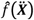
**vi. Back propagate** gradient from label error (i.e. update model, **Eq S18,19**)

**3. Assigning labels to test set** Using test set

**i. Assign cells** to conserved subtypes
**ii. Identify ambiguous label mappings** using cell assignments

##### Algorithm 2 Label Ambiguous Domain Adaption (LAmbDA) Neural Network

**Input:** preprocessed expression matrix 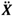, preprocessed labels 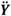, and label mask *G***, Eq S1-3**

**Output:** a trained classifier 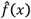 with mapped ambiguous labels and batch effects removed Random initialization

**1. Train on unambiguous labels** Using the subset of samples that have only one possible label For the first half of total iterations:

**i. Forward propagate** predicted labels
**ii. Back propagate** gradient from label error (i.e. update network)

**2. Train on ambiguous labels** Using all samples regardless of number of possible labels For the second half of total iterations:

**i. Forward propagate** predicted labels (i.e. calculate 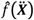, **Eq S14-16**)
**ii. Assign labels** to ambiguously labeled cells (i.e. calculate 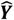, **Eq S8**)
**iii. Calculate Euclidean distances** between subtypes (i.e. calculate *E*, **Eq S9,10**)
**iv. Calculate label error** using 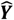 and 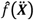
**v. Calculate batch effects error** using ***M*_l_, *M_2_*** and ***E*** (**Eq S10-12**)
**vi. Back propagate** gradient from error terms (i.e. update network, **Eq S20**)

**3. Assigning labels to test set** Using test set

**i. Assign cells** to conserved subtypes
**ii. Identify ambiguous label mappings** using cell assignments

### 2.3 LAmbDA Model Performance

We applied the LAmbDA framework with five different algorithms (LR, FF1, FF3, RNN1, RF) to determine the performance of the LAmbDA-based methods in cell type classification. We measured the following metrics: i) test accuracy of unambiguous labels and corresponding binomial probability of correctly mapping the unambiguous cells (poor mapping = 1.0, accurate mapping = 0.0); ii) cluster-wise distance ratios; iii) Wilcoxon rank sum p-values for comparisons between labels where label ambiguity was added (in the case of pancreas data) and where the true mapping can be inferred in the original publications (in the case of brain data); and iv) a comparison with the MetaNeighbor (Crow, et al., 2018) and CaSTLe packages (Lieberman, et al., 2018).

#### Unambiguous Label Accuracy and Binomial Probability

The test set accuracy of unambiguous labels was generated from the difference between the unambiguous labels and the one-hot predicted labels averaged across each round of cross validation. The weighted accuracy was generated from the mean of each of the individual label accuracies so that each output label was equally weighted. The binomial probability measure was used to calculate the probability of seeing the number of cells correctly assigned to a subtype. Specifically, the binomial probability was the sum of the probabilities that the number of the correctly mapped cells or more would be seen by chance.

#### Distance Ratios to Measure Batch Effects

Three cluster-wise median distance ratios were calculated based on relevant combinations of labels (subtypes) and datasets. The data in these combinations consisted of the Euclidean distances between subtypes of cells in the last hidden layer of the NN implementations of LAmbDA. These combinations were: same dataset-same subtype (*Dat*^+^ *Sub*^+^), which was not used because they were a trivial case that had Euclidean distance = 0.0; same dataset-different subtype (*Dat*^+^ *Sub*^‒^); different dataset-same subtype (*Dat*^−^*Sub*^+^); and different dataset-different subtype (*Dat*^−^*Sub*^−^). For each of the combinations, the median Euclidean distance was calculated from the distances in that group. These median distance values were used to generate 3 ratios for comparison, i) *Dat*^−^*Sub*^+^/*Dat*^+^ *Sub*^−^ (theoretically<1); ii) *Dat*^−^*Sub*^+^/*Dat*^−^*Sub*^−^ (theoretically<1); and iii) *Dat*^+^ *Sub*^−^/*Dat*^−^*Sub*^−^ (theoretically=1, i.e. control). These ratios measured the reduction of dataset batch effects (i), inter-dataset subtype differences (ii), as well as the level of noise introduction by LAmbDA (iii).

#### Assignment of Ambiguous Labels

The label mask (***G*, Supplementary Eq S1**) used in the pancreas datasets had ambiguity added to the label mapping to determine if LAmbDA-FF1 could assign cell types to the correct label. Specifically, possible incorrect label mappings were added to the training mask (***G*, Supplementary Eq S1**). In the brain datasets, we could infer similar mappings between the MusNG and HumN cortical pyramidal cells from past research so we knew the most likely mapping between them (Lake, et al., 2016). These inferred high likelihood mappings were used as further validation. A Wilcoxon rank-sum test was used to measure if LAmbDA-FF1 correctly assigned ambiguous labels to the correct labels in brain or pancreas. Specifically, the number of cells in correct mappings was compared to the number of cells in incorrect mappings using the Wilcoxon rank-sum test. We highlighted the ambiguous label mappings Areas of Interest (AOI) in red, numbered rectangles in the resulting confusion matrices produced by these analyses.

#### Comparison with current methods

We compared LAmbDA-FF1 to CaSTLe and MetaNeighbor. Since CaSTLe could only use two datasets at a time, we used the largest pancreas dataset Bar (8569 cells, 14 labels) to predict the smallest but most diverse dataset Seg (1980 cells, 15 labels). In brain, MusNG (3005 cells, 48 labels) was used to predict HumN (2086 cells, 16 labels). MetaNeighbor predicts the cell label using all of the labels from all datasets. In pancreas this meant 12675 cells across 38 labels and in brain 6376 cells across 70 labels. The unambiguous accuracy was defined as the accuracy during cross validation on the source dataset. The Wilcoxon rank-sum tests were calculated for the same cross dataset comparisons as LAmbDA using weighted accuracy (W-Acc) and area under the curve (AUC)(Bradley, 1997).

## 3 Results

We chose the pancreas datasets to test the feasibility and performances of our methods after introducing ambiguity into the cell type labels, since the pancreas datasets were (i) mostly unambiguous – the labels contained all major cell types with high overlap among all three datasets; (ii) all cells were from the same species and was thus a good testing bed for the label mapping without the added complexity across species. The brain datasets were chosen to test the LAmbDA method capability to deal with issues such as the cross-species complexity, sample imbalance, granularity of labels, and diversity of major cell types. The major cell type classes (e.g. neuron, glial) were labeled in brain too. Therefore we knew the possible subtype mappings in the brain, which served as the ground truth when the performance was evaluated. To evaluate the performance, the batch effects on the unprocessed data had to be analyzed.

The pancreas and brain datasets showed high batch effects, which can be observed from t-SNE diagram (Fig. 1A,B). In this study, LAmbDA aimed at removing the batch effects and revealing conserved subtypes (Fig. 1C) while still maintaining high accuracy in predicting labels of unambiguous cells.

### 3.1 LAmbDA Methods Achieve high accuracy

We compared each of the five LAmbDA-based methods on the pancreas and brain datasets separately. The LAmbDA framework is shown in Fig. 2. All LAmbDA models performed more accurately than random chance (**Supplementary Fig. S3A**, Table 1). The lowest unambiguous accuracy was from LAmbDA-LR in both pancreas data (weighted accuracy: 17%, binomial probability: <1×10^−10^) and brain data (weighted accuracy: 18% binomial probability: <1×10^−10^). The best performing algorithm on unambiguous labels was LAmbDA-RF on both pancreas (weighted accuracy: 91%, binomial probability: <1×10^−10^) and brain data (weighted accuracy: 78%, binomial probability: <1×10^−10^). For mapping ambiguous labels, LAmbDA-FF1 produced the most desirable results (Fig. 3A,C, Fig. 4C,D). LAmbDA-FF1 also maintained high unambiguous accuracy in pancreas data (weighted accuracy: 61% binomial probability: <1×10^−10^) and in brain data (weighted accuracy: 48%, binomial probability: <1×10^−10^, **Supplemenatary Fig. S3A**, Table 1). The LAmbDA-FF1 unambiguous weighted accuracy was similar to that of the more complex LAmbDA-FF3 model (48% vs 49% for pancreas, and 61% vs 67% for brain data). With high unambiguous accuracy, these models were evaluated for their ability to remove batch effects in the data.

**Fig. 2.**
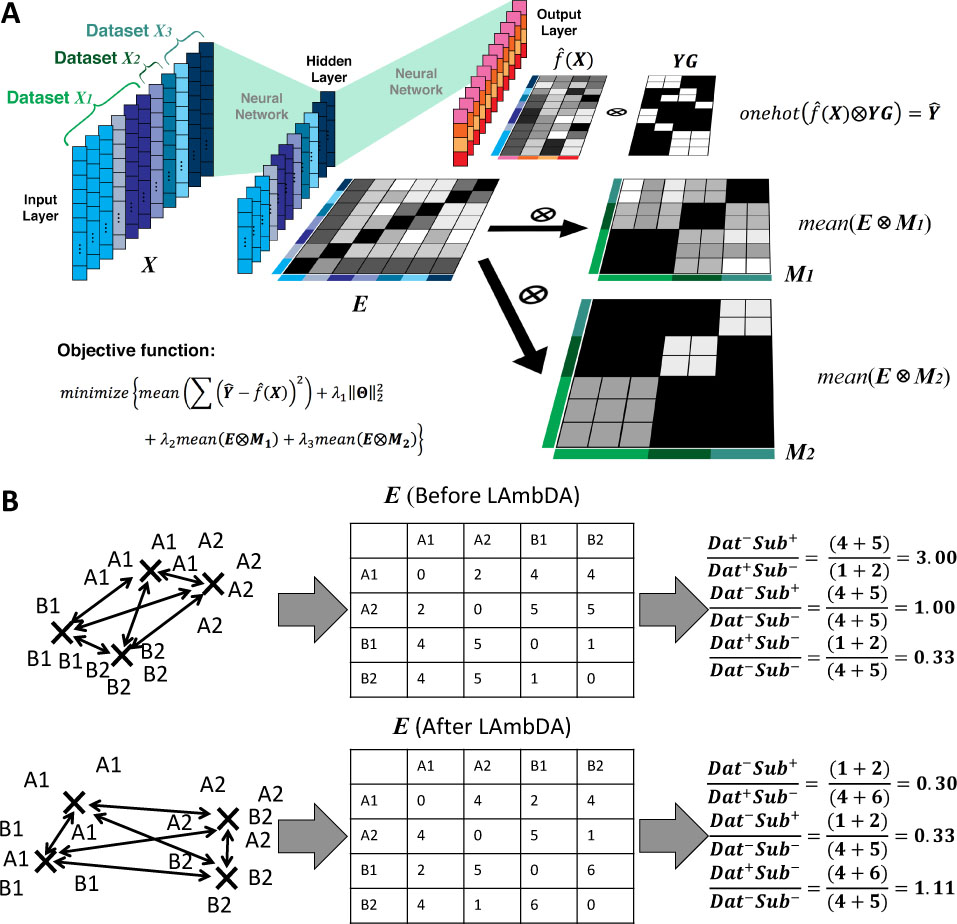
LAmbDA framework: A) The LAmbDA framework including the simplified label mapping (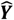, **Eq S8**) and batch effect removal (***E*** ⨂ ***M*_l_**, ***E*** ⨂ ***M_2_***, **Eq S10-12**). B) The distance ratios used to evaluate batch effect reduction where letter indicates dataset and number indicates subtype. The cells are in a reduced feature space in the NN last hidden layer where the distance between subtypes of cells can be measured. The first and second ratio should be less than one and the third ratio should be 1.

**Table 1.** Predictive accuracy and dataset batch effect reduction by LAmbDA model. Full: the full gene set features (i.e. no feature reduction). * indicate that both LR and RF use the full gene set features as input. The cluster distance ratios for LR and RF can be regarded as the full gene set features cluster distance ratios. The LR and RF accuracy can be regarded as the full gene set features accuracy. Distance ratios: i) *Dat*^−^*Sub*^+^ /*Dat*^+^*Sub*^−^, as it approaches 0, subtype increased similarity across datasets. ii) *Dat*^−^*Sub*^+^ /*Dat*^+^*Sub*^−^, as it approaches 0, similar subtypes are closer than dissimilar subtypes. iii) *Dat^−^Sub*^−^ /*Dat^−^Sub*^−^, as it remains near 1, noise is not introduced.

|  | Pancreas |  |  |  | Brain |  |  |  |
| --- | --- | --- | --- | --- | --- | --- | --- | --- |
|  | Distance ratios |  |  | Weighted Accuracy | Distance ratios |  |  | Weighted Accuracy |
|  | i | ii | iii |  | i | ii | iii |  |
| LR | NA* | NA* | NA* | 17% | NA* | NA* | NA* | 18% |
| FF1 | **0.79** | **0.71** | 0.92 | 61% | **0.75** | **0.71** | **0.93** | 48% |
| FF3 | 0.89 | 0.78 | 0.89 | 67% | 1.03 | 0.82 | 0.81 | 49% |
| RNN1 | 0.83 | 0.68 | 0.85 | 31% | 1.32 | 0.71 | 0.55 | 11% |
| RF | NA* | NA* | NA* | **91%** | NA* | NA* | NA* | **78%** |
| Full | 1.12 | 1.04 | **0.95** | NA* | 0.88 | 0.82 | **0.93** | NA* |

### 3.2 LAmbDA Neural Networks Reduce Batch Effects Between Datasets

The neural network-based (NN-based) LAmbDA-FF1, -FF3, and - RNN1 each performed additional feature reduction (Table 1). During training, the hidden layer improved cellular granularity and reduced dataset batch effects as measured by cluster distance ratios (Table 1). LAmbDA-FF1 generated the best reduction of dataset batch effects while still maintaining high cell type signal (Table 1). LAmbDA-FF1 also achieved the best distance ratios overall by reducing the batch effects by 30-32% while introducing 3% noise in pancreas and reducing batch effect distance ratios by 13-15% while only introducing 1% noise in brain (Table 1, **Supplementary Fig. S3B-D**). In the pancreas dataset, LAmbDA-FF1, -FF3, and -RNN1 were able to achieve better distance ratios than the full gene set features (Table 1, **Supplementary Fig. S3B-D).** The brain dataset contained greater batch effects and seemed dependent on the subtype signal. Despite this, LAmbDA-FF1 still outperformed the full feature set across the distance metrics. The datasets themselves showed differing levels of success in batch effect removal.

On relatively simple pancreas datasets, all NN-based models reduced batch effects by 30-35% while only introducing 3-11% noise (Table 1). In more complicated brain datasets, LAmbDA-FF1 was capable of reducing batch effects without introducing noise (Table 1, **Supplementary Fig. S3**). Furthermore, LAmbDA-FF1 correctly learned subtypes that were ambiguously mapped between datasets (Fig. 3A,B **AOI1-3**, Fig. 3C,D **AOI1**).

**Fig. 3.**
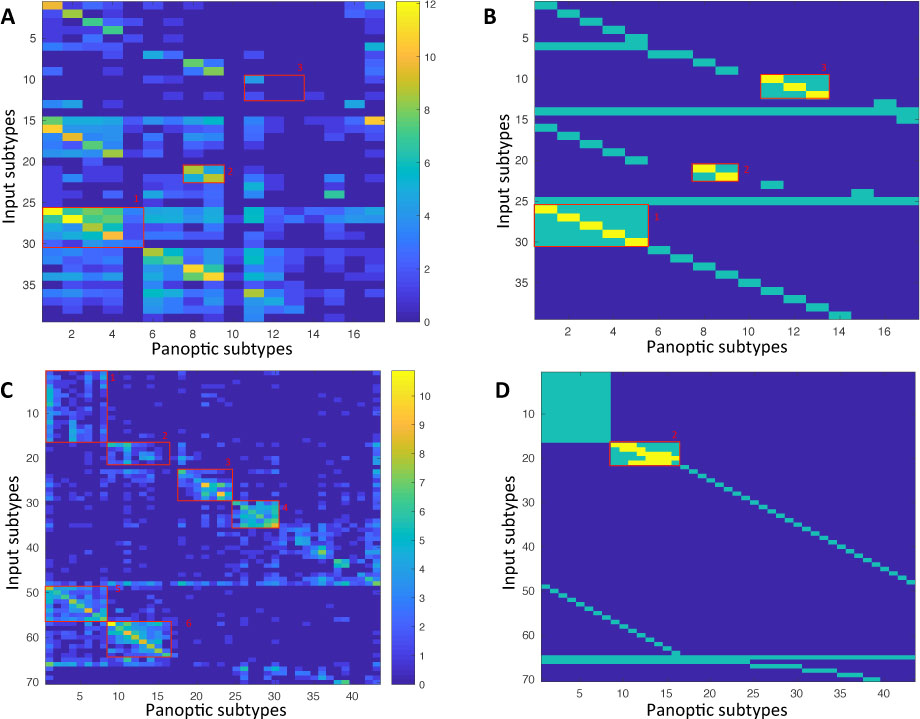
Confusion matrices with their associated label masks used during LAmbDA-FF1 training. Each numbered red box indicates an AOI. A, C) Confusion matrix across three datasets where rows are original cell types and the columns are the conserved cell types (i.e. LAmbDA output labels) for pancreas (A) and brain (C). B, D) The label mask used during LAmbDA training. Green indicates the mask used as input and yellow indicate the true labels, which were either known or inferred from the literature. C) Yellow inside of AOI1-3 indicate true labels from the starting datasets. D) Yellow indicates the cortical layer specific mapping that was inferred from each dataset's publication.

### 3.3 LAmbDA Models Correctly Predict Ambiguous Labels Between Datasets

The LAmbDA-FF1 and LAmbDA-FF3 models correctly mapped pancreatic cells back to their correct label (Wilcoxon p-value: 0.0178 and 0.0346 respectively) when artificial ambiguity was introduced (Fig. 3A,B **AOI1-3**). LAmbDA-FF1 mapped pyramidal cells back to their correct cortical layer (derived from the original papers) across species (Wilcoxon p-value: 0.0181, Fig. 3C,D **AOI2**).

Overall, we found that the general LAmbDA method achieved high accuracy for unambiguous labels regardless which of the five algorithm types were used (LR, FF1, FF3, RNN1, RF). Specifically, if the labels contained low ambiguity, LAmbDA-RF performed most accurately. If there was high ambiguity across datasets, LAmbDA-FF1 performed the most accurately (Table 1). Furthermore the ability to correctly map cortical pyramidal cells shows that cross species comparisons are possible.

### 3.4 High resolution neural subtypes are conserved across species

We discovered that the mouse cortical pyramidal subtypes map to human cortical pyramidal subtypes by their associated cortical layer (e.g. L2 cortex pyramidal cells in mouse are associated with L2 cortex pyramidal cells in human, Fig. 3C **AOI2**, Fig. 3C **AOI1**, Fig. 4D). This indicates that high granularity subtypes are conserved across species (in this case, mouse and human) and the conservation aligns with cortical layer. Because we were able to recreate known or inferred mappings, we applied the mapping from LAmbDA-FF1 interneurons to infer conserved subtypes. These insights allowed us to hypothesize the label mapping of interneurons between human and mouse (Fig. 3C **AOI1**, Fig. 4D). We observed specific subsets of mouse subtypes mapped to the human sub-types. With the biomarkers described in each of the primary sources of the data (Darmanis, et al., 2015; Lake, et al., 2016; Zeisel, et al., 2015), we showed relevant biomarkers for the conserved interneuron subtypes (**Supplementary Table S1**) by intersecting the biomarker lists from the two species. These cross-dataset and -species mappings provided interesting discoveries so we further compared against the two label mapping tools used for scRNA-seq datasets: CaSTLe (Lieberman, et al., 2018) and MetaNeighbor (Butler, et al., 2018).

**Fig. 4.**
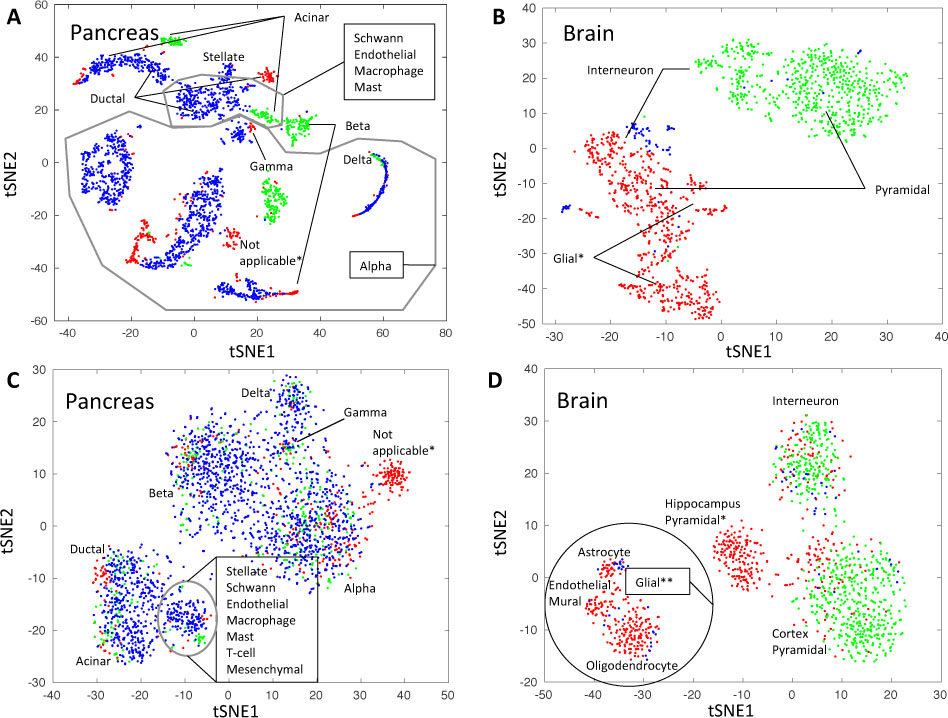
tSNE dimensionality reduction of 20% of samples taken from data before applying LAmbDA (A,B) and after applying LAmbDA (C,D). A) Pancreatic datasets. B) Brain datasets. C) Pancreatic data from the hidden layer of LAmbDA. D) Brain data from the hidden layer of LAmbDA. The colors indicate the dataset A,C) Seger (red), Mur (green), Bar (blue), and B,D) MusNG (red), HumN (green), HumNG (blue). *Indicate cell types that are only present in one dataset. **Indicated glial cells, which are not present in the HumN dataset.

### 3.5 LAmbDA Improves Upon Current Methods

Table 2 describes the performances of LAmbDA, CaSTLe, and MetaNeighbor to predict unambiguous and ambiguous cell types. When the ambiguous labels were tested across datasets, LAmbDA-FF1 had the most significant Wilcoxon p-values indicative of correct mapping (0.0178 and 0.0181). CaSTLe achieved the highest AUC in both pancreas (99%) and brain (94%) data, with LAmbDA-FF1 tied in brain AUC. CaSTLe was also able to achieve the highest weighted accuracy in pancreas (75%). However, these accuracies and AUCs were calculated from the source dataset and could have been caused by overfitting considering the inter-dataset results. Furthermore, the AUC values for both CaSTLe and MetaNeighbor were much closer than the weighted accuracies to LAmbDA-FF1 in all tests. This suggests that CaSTLe and MetaNeighbor are more useful in mapping labels between datasets but should not be used over LAmbDA in classifying individual cells between datasets.

**Table 2.** Performance comparisons between LAmbDA-FF1, CaSTLe, and MetaNeighbor. Pancreas/Brain Map columns contain Wilcoxon rank sum p-values for correct cell labels vs. incorrect cell labels for the groups where artificial label ambiguity was added in LAmbDA. Lower p-values indicate that the algorithm correctly assigned labels between datasets. The Wilcoxon rank-sum p-values were calculated using both the weighted accuracy and AUC. Pancreas/Brain Acc columns contain the weighted accuracy and the mean AUC across all unambiguous labels. The higher the value the better unambiguous labels are fit. In the case of CaSTLe these values were from the source dataset. In MetaNeighbor, these values were from the same dataset and same subtype.

|  | Pancreas Map |  | Brain Map |  | Pancreas Acc |  | Brain Acc |  |
| --- | --- | --- | --- | --- | --- | --- | --- | --- |
|  | W-Acc | AUC | W-Acc | AUC | W-Acc | AUC | W-Acc | AUC |
| LAMBDA-FF1 | 0.0178 | <0.0001 | 0.0181 | 0.0017 | 61% | 94% | 48% | 94% |
| CaSTLe | 0.0632 | 0.0012 | 0.3216 | 0.0038 | 75% | 99% | 32% | 94% |
| MetaNeighbor | 0.7446 | <0.0001 | NaN | 0.0041 | 53% | 86% | 20% | 86% |

### 3.6 Major Cell Types Consistent Across Species and Dataset

Aside from the mapping of ambiguous labels across datasets, we found consistent mapping patterns between subtypes within the same major cell type. These mappings further validate our method. For example, the MusNG oligodendrocyte subtypes showed high consistency with other oligodendrocyte subtypes compared to other subtypes (Wilcoxon p-value = 1.67×10^−30^, Fig. 3C **AOI4**, Fig. 4D). The HumNG oligodendrocytes mapped to multiple MusNG oligodendrocytes compared to other subtypes (Wilcoxon p-value = 1.51×10^−3^, Fig. 4D), and the HumNG astrocytes mapped to multiple MusNG astrocyte subtypes compared to other subtypes (Wilcoxon p-value = 1.62×10^−5^, Fig. 4D).

Cortical interneuron subtypes were highly consistent with other cortical interneuron subtypes in HumN compared to other subtypes (Wilcoxon p-value = 5.17×10^−48^, Fig. 3C AOI5, Fig. 4D), and cortical pyramidal subtypes were highly consistent with other cortical pyramidal subtypes in HumN compared to other subtypes (Wilcoxon p-value = 3.94×10^−35^, Fig. 3C AOI6, Fig. 4D). Such relationships were observed in the pancreas data, where immune cells clustered with one another (Fig. 4C). Furthermore, we found that models trained with MusNG and tested on HumN and vice versa showed the same major cell type patterns (**Supplementary Fig. S2**).

## 4 Discussion

All LAmbDA-based methods improved the prediction of unambiguous cell type accuracy between datasets, with each LAmbDA model catering to different specific demands. For instance, LAmbDA-FF1 performs best at correctly removing batch effects. LAmbDA-RF is most accurate at predicting unambiguous labels. LAmbDA-RNN1 shows desirable characteristics in integrating the datasets, but needs to be further optimized. We suggest different LAmbDA models should be considered to suit different dataset ambiguity levels. These considerations are especially important when studying the correct assignment of ambiguous labels.

We observed that when error is intentionally introduced into the labels, LAmbDA models were still able to correctly identify the labels in pancreas and brain tissue (artificial ambiguity 10 in 39 labels in pancreas and 5 in 70 labels in brain). These errors were introduced when the label mappings were known but were not included. LAmbDA can identify the correct label in most cases (Fig. 3B **AOI1-3**, Fig. 3D **AOI1**). This is in part due to the feature reduction step in the NN implementations which rearrange the subtype clusters to reduce batch effects. Even after feature reduction, we see interesting subtype mappings both within and between datasets/species.

Similar subtypes within a species tend to cluster together. For instance, in the brain, the oligodendrocyte cell types in MusNG formed a consistent group. This implies that subtypes of cells are difficult to further stratify and consist of a joint distribution of major cell types within the brain layer. Mouse and human interneurons from the LAmbDA-FF1 model were mapped to each other. They can be considered conserved subtypes, which are consistent across dataset and species. We used the intersection of biomarkers from the previous publications to identify these conserved subtypes.

An interesting cell mapping pattern was the HumNG subtypes tended to map to the MusNG subtypes more often than HumN, especially before batch effect removal in the full feature set. One possible reason is that HumN was single nuclei sequencing as opposed to whole cell sequencing in HumNG and MusNG, so the gene expression profiling could be quite different. This suggests that sequencing method may introduce larger batch effects than species differences, and cross-species training of models may be more feasible than once thought. Due to these considerations we believe that the general LAmbDA framework has a great deal of potential.

These applications of LAmbDA-based models on brain and pancreas data make compelling cases for the LAmbDA method. We postulate that our method can also adopt other learning algorithms such as deep learning as well as other distance metrics for the hidden layer to improve its dataset/species integration and prediction accuracy. We also believe that the LAmbDA framework is model-independent because of the high accuracy and batch effect removal achieved by multiple tested models, thus making it ideal for incorporation with other machine learning models. Furthermore, even though scRNA-seq data was used in our study, the LAmbDA framework is not fundamentally limited to any data type, organism, or disease. For instance, disparate tumor datasets could be combined to find conserved cell populations between patients, datasets, and similar cancer types (e.g. grades of glioma).

The scalability of LAmbDA is immense. Since LAmbDA does not compute any pairwise correlations between samples, it could be easily scaled up to incorporate the increasing number of large Drop-seq datasets for single-cell studies. It is also worth mentioning that the core of the LAmbDA framework is a set of cost functions in Python (TensorFlow), making it ideal for others to integrate into their own workflows.

## 5 Conclusion

We developed a novel dataset integration and ambiguous subtype labeling framework, LAmbDA, to predict cellular subtypes. Our algorithm addresses both label mapping and dataset batch effect issues simultaneously. We are able to perform these analyses without exact label correspondence. Our method is ideal to scale to even larger datasets. LAmbDA proves to be accurate for subtype prediction across species and datasets. It is model independent and capable of revealing hidden biological relationships between subtypes in disparate datasets. This could prove especially useful in identifying conserved cell populations across tumors or stages. Furthermore, in theory, this method could be applied to any scalar data, which contain multiple datasets and ambiguous label mappings. LAmbDA can be integrated into existing machine learning pipelines to identify conserved labels and improve the robustness of the model to data systematic biases.

## Acknowledgements

The authors thank the faculty and students at the Indiana University Purdue University Indianapolis School of Informatics and Computing and Center for Computational Biology and Bioinformatics for their input and technical expertise.

