## Supplementary Materials for "LAmbDA: Label Ambiguous Domain Adaption Dataset Integration Reduces Batch Effects and Improves Subtype Detection"

### Supplementary Material

#### 1.1) Comparison of major neural cell types

##### Methods

First, we wanted to establish the viability of cross-species comparisons at the major cell type level for neurons and glia. We used the MusNG and HumN datasets. We identified a 439 gene feature set using maximum relevance minimum redundancy (Ding and Peng, 2005) in mouse to differentiate the major cell types located in the brain (interneuron, S1 pyramidal, CA1 pyramidal, oligodendrocyte, microglia, endothelial, astrocyte, ependymal, mural, as identified by the Linnarson group (Zeisel, et al., 2015)). ependymal and mural cell types were not identified in the HumN dataset and therefore were used as a negative control.

Before the data was submitted to mRMR, we performed feature selection (Zeisel, et al., 2015) consisting of i) removing low expression genes, ii) removing genes that do not coexpress with other genes and iii) retaining the top 5000 high variance genes using a noise model fit to the data (Zeisel, et al., 2015). The samples were separated into the 9 major cell types with the 5000 highest variance genes. This labeled dataset was submitted to the mRMR algorithm resulting in 500 optimally orthogonal and high variance genes. After filtering genes that matched between the MusNG and HumNG datasets, the 500 mRMR-selected genes were further reduced to 439.

These genes were then used to train a NN classifier on the MusNG data separated into 3 groups, train (60%), validation (20%) and test (20%). The model was trained using 10 $L_2$ regularization values (Lambda: 0.0, 0.001, 0.01, 0.05, 0.2, 0.5, 1, 2, 5 10, Fig 4) on the training set, and the final lambda value was chosen by selecting the lambda value with the highest overall accuracy on the validation set. The resulting highest accuracy NN model was used to predict cell types in the MusNG test set and in the entire HumNG dataset (Darmanis, et al., 2015).

As a comparison, a regularized logistic regression (LR) classifier was trained on each of the cell types using the same 439 gene feature set. The training set consisted of the same samples as the NN classifier. The validation set and test set were also the same NN samples. LR was conducted using the same lambda values as the NN classifier resulting 90 models, 9 for each cell type for 10 different lambda values. The best performing lambda (across all cell types) selected by overall accuracy on the test set was used for all 9 models and evaluated on the MusNG validation set and the entire HumNG dataset. We evaluated the accuracy of these models using overall accuracy $\left(\frac{TP}{(TP+FP+TN+FN)}\right)$ and weighted accuracy $\left(\sum_{i=1}^{|\{major\ cell\ types\}|}\frac{TP_i}{|\{major\ cell\ types\}|\times(TP_i+FP_i+TN_i+FN_i)}\right)$ across each cell type ($i \epsilon\ major\ cell\ types$).

##### Results

We observed that the regularized neural network (NN) model performed better than the regularized logistic regression (LR) model in the mouse dataset (overall accuracy: LR = 0.87 and NN = 0.91 respectively, **Supplementary Fig. S1**). Moreover, the overall accuracy increased greatly when comparing across species (overall accuracy: LR = 0.4 and NN = 0.55, **Supplementary Fig. S1**). We also see the negative controls (ependymal and mural, I.e. no correspondence in the human dataset) have few cells assigned to them when using the NN (**Supplementary Fig. S1D**).

Aside from the overall accuracy, which was biased by the number of cells in each cell type, we wanted to view the weighted accuracy to get a representation of the performance across cell types. We found that the NN classifier performed better in the mouse validation set (weighted accuracy: LR = 0.69 and NN = 0.74, **Supplementary Fig. 1**). More intriguingly, the human dataset weighted accuracy was even more improved (weighted accuracy: LR = 0.46 and NN = 0.61, **Supplementary Fig. S1**).

These results showed that the gene feature set was useful between species but could be improved to provide more accurate classification. There are clear advantages of using *a priori* feature sets to identify cell type similarity (Crow, et al., 2018) but we also aim to explore how feature reduction through a neural network could identify corresponding neural subtypes.

#### 1.2) Comparison of neural subtypes

#### Methods

Next, we wanted to explore the more granular interneuron and pyramidal subtypes available to us in HumN and MusNG datasets. The mouse dataset constituted both neural and glial cell types while the human dataset constituted only neural cell types. This lack of cell types in human provided us with a negative control. We needed to retain more features due to the increased granularity of the subtypes (Supplementary Material Sec 1.2) vs. major cell types (Supplementary Material Sec 1.1). First, we identified the overlapping gene feature set between both datasets (13355 genes). Next, we converted the cell subtypes into numeric form so that a model could be trained. Because the NN model performed better than a logistic regression classifier in our previous experiment, we only trained NN classifiers. We performed 50 iterations random cross validation on mouse NN classifiers (80% training and 20% testing) using all 48 cell type labels. This process was repeated for multiple lambda regularization values (0.0, 0.001, 0.01, 0.1, 0.5, 1, 1.5, 2, 2.5, 3, 3.5, 4, 4.5, 5). These multiple classifiers corresponded to the multiple lambda regularization values that we wanted to test. This model then was used classify MusNG test set and entire HumN dataset.

Because the major cell types are known in human and mouse, we recorded the accuracy in identifying the major cell types. The same process was repeated for the human dataset in which a model was trained on 80% of the human dataset and tested on 20% 50 times. Because the human dataset only contained neural cell types, only mouse neuron subtypes were used in the human-trained NN classifier. For both datasets, the major type accuracy was recorded for both mouse and human test sets. The subtype level accuracy was recorded for the within species test set.

#### Results

We observed that the major divisions of interneurons and pyramidal cells were conserved across species. The mouse trained model achieved mouse subtype accuracy of 67% and mouse major cell type accuracy of 93% (**Supplementary Fig. S2A,C**). The mouse trained model was also able to accurately identify major cell types in human cells (**Supplementary Fig. S2A,C**): i) human interneurons were classified interneuron (anova p-value=$8.17\times10^{-9}$), ii) human pyramidal cells were classified pyramidal (anova p-value=$4.87\times10^{-6}$), iii) human cells classified interneuron were interneuron (anova p-value=$1.06\times10^{-6}$), and iv) human cells classified pyramidal were pyramidal cells (anova p-value=$1.14\times10^{-2}$). We saw similarly high levels of accuracy in human.

The human trained model achieved human subtype accuracy of 94% and human major cell type accuracy of 99% (**Supplementary Fig. S2B,D**, note: the human dataset contained fewer subtypes and major cell types increasing the accuracy). The human trained model was also able to accurately identify major cell types in mouse cells (**Supplementary Fig. S2B,D**): i) mouse interneurons were classified interneuron (anova p-value=$6.00\times10^{-11}$), ii) mouse pyramidal cells were classified pyramidal (anova p-value=$1.55\times10^{-2}$), iii) mouse cells classified interneuron were interneuron (anova p-value=$3.75\times10^{-4}$), and iv) mouse cells classified pyramidal were pyramidal cells (anova p-value=$3.38\times10^{-3}$).

In human, the cell types were rarely associated with a single mouse cell type but rather related to combinations of mouse subtypes. For example, human cell type: In6(6) was related to mouse cell type: Int2(2) and Int14(14) in both mouse and human trained models. Human cell type: In1(1) was related to mouse cell type: Int12(12) (Fig 4). Aside from the interneuron cells, we saw relationships in the pyramidal cell types as well. Human cell type: Ex1(9) was related to mouse cell type: S1PyrL23(17) in both human and mouse trained models. Human cell type: Ex3(11) was related to mouse cell type: S1PyrL4(18) in both human and mouse trained models. These relationships advance the idea that these tissue specific subtype profiles are conserved across species and can be leveraged to identify conserved neural subtypes (**Supplementary Fig. S2**).

Also, we found that the cortical layer specific subtypes were mirrored in the classification between species. This can most prominently be seen in **Supplementary Fig. S2C** where human and mouse cells from the same cortical layers correspond to one another (i.e. Ex1(9)→S1PyrL23(17), Ex2(10)→S1PyrL4(18), Ex3(11)→S1PyrL4(18), Ex4(12)→S1PyrL4(18), Ex5(13)→S1PyrL5(19), Ex6→S1PyrL5(19)/S1PyrL6(20)).

#### 2.1) Detailed Implementation

##### Data Preprocessing

To follow the algorithms above, we convert the general framework above into a concrete implementation. The possible label mappings of the original subtypes to the conserved subtypes are established from the literature (Darmanis, et al., 2015; Lake, et al., 2016; Zeisel, et al., 2015). We determine the known relationships between $L$ and $\hat{L}$ and represent them using the label mask ($\boldsymbol{G}$ in **Eq S1**). A subtype $L_i$ can be exactly the same subtype as $\hat{L}_j$ (e.g. $MusNG\ Interneuron\ 5\ \equiv MusNG\ Interneuron\ 5$ ), potentially the same subtype as $\hat{L}_j$ (e.g. $MusNG\ Interneuron\ 5\ \cong HumN\ Interneuron\ 2$ ), or not the same subtype as $\hat{L}_j$ (e.g. $MusNG\ Interneuron\ 5\ \neq HumNG\ Oligodendrocyte$).

$$\boldsymbol{G} \in \mathbb{Z}^{l \times \hat{l}}$$

$$\boldsymbol{G_{i,j}} = \begin{cases} 1, & L_i \equiv \widehat{L_j} \\ 1, & L_i \cong \widehat{L_j} \\ 0, & L_i \neq \widehat{L_j} \end{cases} \qquad \text{Eq S1}$$

Once the proper label mask ($\boldsymbol{G}$) is generated (**Eq S1**) all of the necessary data is available to run LAmbDA. These data must now be processed before performing Algorithm 1 or 2. Firstly, only genes with variance in the top $20^{th}$ percentile and with fewer than 50% zeros are selected. The entire matrix is divided by 10 and log2 transformed ($log2(\boldsymbol{X}/10 + 1)$). Next, sample imbalances are addressed by over-sampling under-represented cell subtypes and under-sampling the over-represented cell subtypes. Note that our testing set during cross-validation is removed at this point. A cutoff ($c$, **Eq S2**) is selected based on a percentile of the label distribution for over and under sampling. The quartile is selected using the optunity (Claesen, et al., 2014) hyperparameter search tool. Optunity searches distributions of hyperparameters based on a cost function provided by the user and returns the set of hyperparameters that minimize the cost function. The resampling is also weighted by the ambiguity of the subtypes (i.e. number of possible labels for a subtype) using a constant ($\gamma$). Note that if $\gamma$ is set to 0 then there will be no weighting based on the label ambiguity.

$$c = percentile(colSums(\boldsymbol{Y}), p_{cut})$$

$$o = c \left\lceil \log_2 \left( \left( \frac{\max(rowSums(\boldsymbol{G}))}{rowSums(\boldsymbol{G})} \right) + 1 \right)^{\gamma} \right\rceil \qquad \text{Eq S2}$$

Variable $A$ is the set of each individual row in $\boldsymbol{Y}$. The combined over/under-sampling t cutoff point ($o$) results in a new set of indices $\ddot{A}$. The new set of sample indices ($\ddot{A}$) were then used to index our original data matrix ($\boldsymbol{X}$) and label matrix ($\boldsymbol{Y}$) to produce our final data matrix ($\ddot{\boldsymbol{X}}$) and our final label matrix ($\ddot{\boldsymbol{Y}}$, **Eq S3**).

$$\ddot{\boldsymbol{X}} = \boldsymbol{X}_{(:),\ddot{A}}\,, \quad \ddot{\boldsymbol{Y}} = \boldsymbol{Y}_{(:),\ddot{A}},\ \ddot{n} = |\ddot{A}| \qquad \text{Eq S3}$$

#### Assigning labels to ambiguously labeled cells

Due to systematic bias in the datasets, we introduce a dispersion constant ($\tau$) to prevent subtypes from being selected in early iterations and reinforced at a local minima. The dispersion constant is selected using the optunity (Claesen, et al., 2014). To prevent any weighting, the dispersion constant can be set to 0. The dispersion-weighted ($d$) matrix is denoted $\hat{f}(\ddot{\boldsymbol{X}})^{(d)}$ (**Eq S4**). The constant of ε (=0.1) is added to prevent artificial masking of desired labels.

$$\hat{f}(\ddot{\boldsymbol{X}})^{(d)}_{(i),j,:} = \left(\hat{f}(\ddot{\boldsymbol{X}}_{(i)}) + \varepsilon\right)_{(i),j,:} \left[\frac{mean(\hat{f}(\ddot{X}_{(i)})+\varepsilon)}{colMeans(\hat{f}(\ddot{X}_{(i)})+\varepsilon)}\right]^{\tau} \qquad \text{Eq S4}$$

$$|\ i,j \in \mathbb{Z}, 1 \leq i \leq 3, 1 \leq j \leq \ddot{n}$$

Then, the label mask ($\boldsymbol{G}$) is used to set all incorrect labels to 0 so that they will not be selected as the correct label. The masked ($m$) dispersion-weighted ($d$) matrix is denoted $\hat{f}(\ddot{\boldsymbol{X}})^{(m)(d)}$ (**Eq S5**).

$$\hat{f}(\ddot{\boldsymbol{X}})^{(m)(d)} = \ddot{\boldsymbol{Y}}\boldsymbol{G} \otimes \hat{f}(\ddot{\boldsymbol{X}})^{(d)} \qquad \text{Eq S5}$$

Each of the original subtype labels in matrix $L$ should be grouped with similar subtypes $\hat{L}$. To enforce this, a subtype weight is added, resulting in the subtype weight matrix ($\hat{f}(\ddot{\boldsymbol{X}})^{(s)}$) (**Eq S6**).

$$\hat{f}(\ddot{\boldsymbol{X}})^{(s)} = \ddot{\boldsymbol{Y}}\ddot{\boldsymbol{Y}}^{\mathrm{T}}\hat{f}(\ddot{\boldsymbol{X}})^{(m)(d)} \qquad \text{Eq S6}$$

The subtype label weights then are combined with the masked dispersion weighted matrix at a proportion specified by Δ. To prevent any correction to the original subtype weights, Δ can be set to 0 (**Eq S7**).

$$\hat{f}(\ddot{\boldsymbol{X}})^{(s)(m)(d)} = \Delta\hat{f}(\ddot{\boldsymbol{X}})^{(s)} + (1-\Delta)\hat{f}(\ddot{\boldsymbol{X}})^{(m)(d)} \quad \text{Eq S7}$$

The subtype-weighted masked dispersion weighted output ($\hat{f}(\ddot{\boldsymbol{X}})^{(s)(m)(d)}$) now can be converted into labels using one-hot encoding shown below. It is important to note the labels $\hat{L}$ that have a direct correspondence in $L$ can only be assigned to one member of $\hat{L}$. These labels with only one possible mapping are called unambiguous and used in later analyses to determine algorithm performance.

$$\widehat{\boldsymbol{Y}} = onehot\left(argmax\left(\hat{f}(\ddot{\boldsymbol{X}})^{(s)(m)(d)}\right)\right) \quad \text{Eq S8}$$

Once **Eq S8** is calculated, only function definitions ($\hat{f}(\ddot{\boldsymbol{X}})$) and the loss functions are needed to perform Algorithm 1 for LR and RF. For the NN-based models, the batch effects are removed in the final hidden layer before the subtype predictions are made.

#### Calculating Batch Effect Error

To remove the batch effects introduced by using multiple datasets we use Euclidean distance between subtypes to reduce inter-dataset distances. In this section, a one hidden layer example is used but can be changed to fit other NN architectures by adding additional layers ($H_{(i)}$). The hidden layer ($\boldsymbol{H}$) is a regularized dropout layer with sigmoid activation function and $n_{hidden}$ hidden nodes. The drop out percentage ($p_{drop}$) is optimized using sobol search from the optunity package (Claesen, et al., 2014). Dropout is used in the hidden layer(s) because it has been shown to be more robust in high random noise datasets (Rodner, et al., 2016) like scRNA-seq data. To reduce the dataset biases, we add the loss term masks $\boldsymbol{M_1}$ (**Eq S11**), and to increase differences within datasets between labels we add $\boldsymbol{M_2}$ (**Eq S12**). These new constraints are added to the hidden layer outputs using ambiguous label centroid ($\boldsymbol{C}$ in **Eq S9**) squared Euclidean distances ($\boldsymbol{E}$ in **Eq S10**) inspired by computer vision research (Wen, et al., 2016). Note that "tril" denotes the lower triangle of a matrix. With the processing steps (i.e. ambiguous label assignment and batch effects removal) defined, the function ($\hat{f}(x)$) for each model type can be defined.

$$\ddot{\boldsymbol{Y}}_{\boldsymbol{colsum}} = \begin{bmatrix} colSums(\ddot{\boldsymbol{Y}})_1 \\ \vdots \\ colSums(\ddot{\boldsymbol{Y}})_{\ddot{n}} \end{bmatrix}$$

$$\boldsymbol{C} = sigmoid(\ddot{\boldsymbol{X}}\boldsymbol{H})\ddot{\boldsymbol{Y}} \oslash \ddot{\boldsymbol{Y}}_{\boldsymbol{colsum}} \quad \text{Eq S9}$$

$$\boldsymbol{E} = rowSums(\boldsymbol{C}^2)1^{1\times l} + 1^{l\times 1}rowSums(\boldsymbol{C}^2)^T - 2\boldsymbol{C}\boldsymbol{C}^{\boldsymbol{T}} \quad \text{Eq S10}$$

$$\boldsymbol{M_1} = \frac{(\boldsymbol{G}\boldsymbol{G}^{\boldsymbol{T}}>0)\otimes(1^{l\times l}-(\boldsymbol{D}\boldsymbol{D}^{\boldsymbol{T}}>0))}{tril\left(rowSums(\boldsymbol{G})1^{1\times l}\right)+tril\left(rowSums(\boldsymbol{G})1^{1\times l}\right)^T} \quad \text{Eq S11}$$

$$\boldsymbol{M_2} = \frac{(\boldsymbol{D}\boldsymbol{D}^{\boldsymbol{T}}>0)}{tril\Big(rowSums\big((\boldsymbol{D}\boldsymbol{D}^{\boldsymbol{T}}>0)\big)\Big)+tril\Big(rowSums\big((\boldsymbol{D}\boldsymbol{D}^{\boldsymbol{T}}>0)\big)\Big)^T} \quad \text{Eq S12}$$

#### Defining Functions for Each Model Type

We study five different machine learning functions ($\hat{f}(x)$) on the processed scRNA-seq data ($\ddot{\boldsymbol{X}}$). Specifically, $\hat{f}(\ddot{\boldsymbol{X}})$ is calculated for LR (**Eq S13**), FF1 (**Eq S14**), FF3 (**Eq S15**), RNN1 (**Eq S16**), and RF (**Eq S17**). The output layer ($\boldsymbol{O}$) is regularized with a softmax activation function. The resulting label matrix $\widehat{\boldsymbol{Y}}$ is used to train an individual iteration of our model ($\hat{f}(\ddot{\boldsymbol{X}})$). The process allows for ambiguous labels to change iteratively to another possible label (**Fig 2B**). The resulting process assigns these ambiguously labeled cells to refined subtypes. For FF3 **Eq S15** instead has three different $\boldsymbol{H}$ variables nested in the equation symbolizing the three layers (**Eq S16**). Similarly, the recurrent NN implementation has a recurrent layer ($\boldsymbol{R}$) instead of a single $\boldsymbol{H}$ layer (**Eq S17**). The RF method uses aggregate probabilities from trees ($\hat{t}(x)$) with the same scRNA-seq data ($\ddot{\boldsymbol{X}}$). With the models defined, the loss terms to be optimized can be described in detail.

$$\hat{f}(\ddot{\boldsymbol{X}}) = softmax(\ddot{\boldsymbol{X}}\boldsymbol{O}) \quad \text{Eq S13}$$

$$\hat{f}(\ddot{\boldsymbol{X}}) = softmax(sigmoid(\ddot{\boldsymbol{X}}\boldsymbol{H})\boldsymbol{O}) \quad \text{Eq S14}$$

$$\hat{f}(\ddot{\boldsymbol{X}}) = softmax(sigmoid(sigmoid(sigmoid(\ddot{\boldsymbol{X}}\boldsymbol{H_1})\boldsymbol{H_2})\boldsymbol{H_3})\boldsymbol{O}) \quad \text{Eq S15}$$

$$\hat{f}(\ddot{\boldsymbol{X}}) = softmax(sigmoid(\ddot{\boldsymbol{X}}\boldsymbol{R})\boldsymbol{O}) \quad \text{Eq S16}$$

$$\hat{f}(\ddot{\boldsymbol{X}}) = \frac{1}{n_{trees}}\sum_{i=1}^{n_{trees}}(\hat{t}(\ddot{\boldsymbol{X}})) \quad \text{Eq S17}$$

##### Defining Cost Function for Each Model Type

The cost functions for different model types were slightly different based on the model characteristics most specifically the presence of hidden layer(s). Note the sum of $H_{(i)}$ denotes the sum of all weights on all layers. The RF model used the cost function in **Eq S18** due to the lack of an L2 regularization term. The LR model used the cost function in **Eq S19** since there was no hidden layer but still an L2 regularization term. The three types of NN implementations of LAmbDA (FF1, FF3, and RNN1) (Aymeric, 2018) use the cost function in **Eq S20**. Once the cost functions are defined the model can be trained using most machine learning frameworks like TensorFlow.

$$\min\left(mean\left(\sum\left(\widehat{\boldsymbol{Y}} - \hat{f}(\ddot{\boldsymbol{X}})\right)^2\right)\right) \quad \text{Eq S18}$$

$$\min\left(mean\left(\sum\left(\widehat{\boldsymbol{Y}} - \hat{f}(\ddot{\boldsymbol{X}})\right)^2\right) + \lambda_1\sum\sum(\boldsymbol{H}_{(i)}{}^2)\right) \quad \text{Eq S19}$$

$$\min\left(mean\left(\sum\left(\widehat{\boldsymbol{Y}} - \hat{f}(\ddot{\boldsymbol{X}})\right)^2\right) + \lambda_1\sum\sum(\boldsymbol{H}_{(i)}{}^2) + \lambda_2 mean(\boldsymbol{E}\otimes\boldsymbol{M_1}) - \lambda_3 mean(\boldsymbol{E}\otimes\boldsymbol{M_2})\right) \text{Eq S20}$$

##### Additional Information for Running the Algorithms

To train the LAmbDA models, we use the AdamOptimizer (Kingma and Ba, 2014) with step size of 0.01 and random mini-batches of size $p_{batch}$ (a percentage of $\ddot{n}$) that were changed every 50 iterations to prevent overfitting the unambiguous labels. We ran each model for 2000 iterations except for the Random Forest model, which was run for 100 iterations. The code is written for GPU-enabled TensorFlow Python3 package. There were some other considerations that needed to be accounted for during training relating to specific models.

Because recurrent NNs use data order in training, the graph structure within the RNA co-expression data is used to sort the genes data matrix ($\boldsymbol{X}$). Using the top 5000 genes with highest variance, the principle components (PCs) are calculated and then the genes are sorted by the first PC. Next, the expression of the top 5000 gene values are sorted into an 100 by 50 matrix before training, where the sorted genes are separated into 100 rows with 50 genes in each row. This procedure allows for a part of the co-expression network structure to be utilized by the RNN.

There are two non-NN based models tested which do not contain hidden layers: LR and RF. LAmbDA-LR uses the same hyperparameters as the NN models except that $n_{hidden}$, $\lambda_2$, and $\lambda_3$ are not needed due to a lack of a hidden layer. LAmbDA-RF does not contain $n_{hidden}$, $\lambda_1$, $\lambda_2$, and $\lambda_3$ but requires other hyperparameters related to the trees in the random forest. These features are number of trees in the forest ($n_{trees}$) and maximum number of nodes in the trees ($n_{max}$). In both of these circumstances the necessary hyperparameters are tuned using the sobol solver in the optunity package (Claesen, et al., 2014).

#### 2.2) LAmbDA model output

Figures 3 and 4 in the main text were also generated for the other LAmbDA variants to compare and contrast the results. The best overall results were using the LAmbDA-FF1 variant and so those figures are included in the main text. The other variants in general had good unambiguous accuracy but were not as high accuracy as LAmbDA-FF1. One important feature of note is that the LAmbDA-RNN1 model does have good characteristics in integrating datasets evenly despite the accuracy drawbacks (**Supplementary Fig. S9).** The LAmbDA-RF variant has high unambiguous accuracy but does not transfer the knowledge well between datasets (**Supplementary Fig. S7**).

#### 2.3) LAmbDA Hyperparameter Optimization

##### Methods

Multiple types of models were trained using the LAmbDA method, which required the tuning of multiple hyperparameters. We tuned these parameters using a sobol solver in the optunity package (Claesen, et al.,

2014). During the training, we output these parameters into a file so that the parameters could be studied afterwards. Each algorithm was trained 50 times using the sobol-distributed hyperparameters for each iteration of cross validation. The result is that for each of those 500 trained models we have the error and all of the hyperparameter settings. With these we plot the hyperparameters against the error to better understand how they affect model training.

#### Results

Some of the hyperparameters were more associated with the training error than others. For instance, increased $\tau$ tended to increase the error during training (**Supplementary Fig. S10**). However if $\tau$ is set too low then many of the subtypes may not be able to disperse properly. Also, higher dropout rate also tended to be associated with higher error. The higher the percentage cutoff also tended to be associated with lower error. Higher $\gamma$ also tended to be associated with lower error. Though this is seen more drastically in the brain dataset. Another feature of note is that the brain and pancreas dataset though in general mirroring each other's patterns were not identical. This could be due to the higher granularity of the brain dataset.

### Supplementary Figures

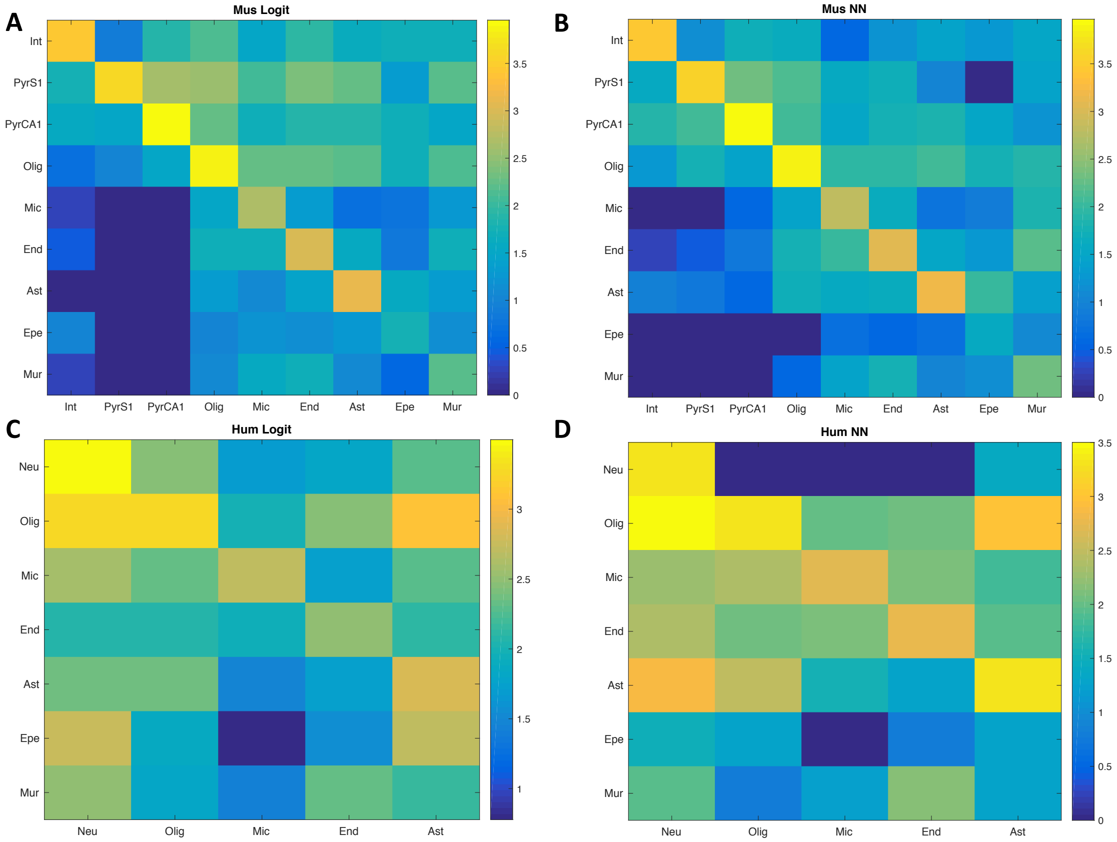


**Supplementary Fig. S1.** Logit corresponds to the logistic regression model. NN corresponds to the neural network model. For each confusion matrix the rows correspond to predicted major cell types (MusNG) and the columns correspond to true cell types (either MusNG or HumNG). A) The Logit classification confusion matrix (log2) on the MusNG test set (rows: MusNG preds, cols: MusNG labels). B) The NN classification confusion matrix (log2) on the MusNG test set (rows: MusNG preds, cols: MusNG labels). C) The Logit classification confusion matrix (log2) on the HumNG dataset (rows: MusNG preds, cols: HumNG labels). D) The NN classification confusion matrix (log2) on the HumNG dataset (rows: MusNG preds, cols: HumNG labels). Row/Col labels: Neu (neuron), Olig (oligodendrocyte), Mic (microglia), End (endothelial), Ast (astrocyte), Epe (ependymal), and Mur (mural).

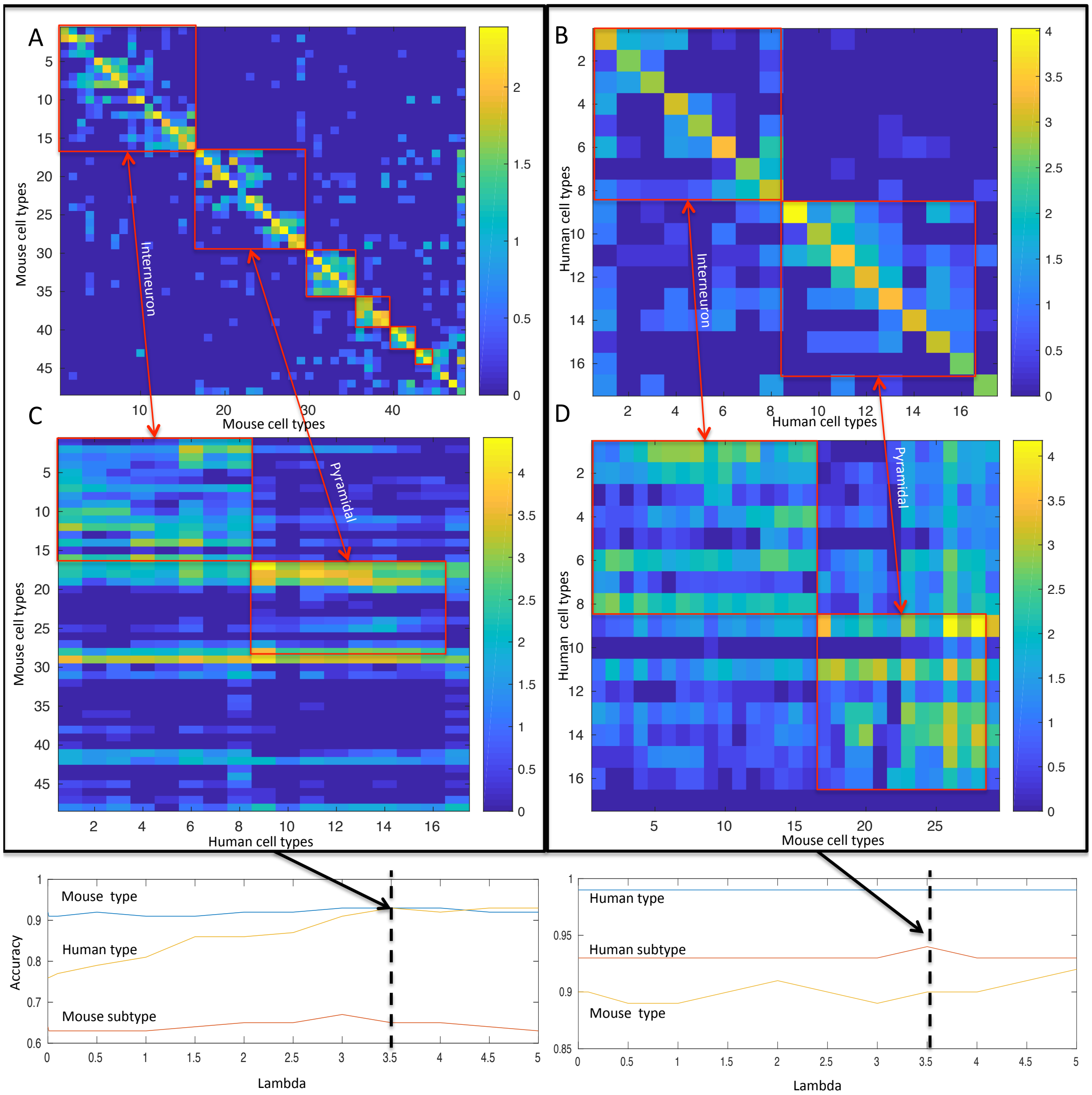


**Supplementary Fig. S2.** Red boxes indicate major cell types. The two largest red boxes in each panel indicate either interneurons or pyramidal cells. A) The confusion matrix (log2) of the MusNG trained model on the MusNG test set over 50 fold cross valdation at lambda of 3.5. B) The confusion matrix (log2) of the HumN trained model on the HumN test set over 50 fold cross validation at lambda of 3.5. C) The confusion matrix (log2) of the MusNG trained model on the HumN dataset over 50 fold cross validation. D) The confusion matrix (log2) of the HumN trained model on the MusNG dataset over 50 fold cross validation. The bottom panel shows the change in major cell type and subtype accuracy with change in lambda.

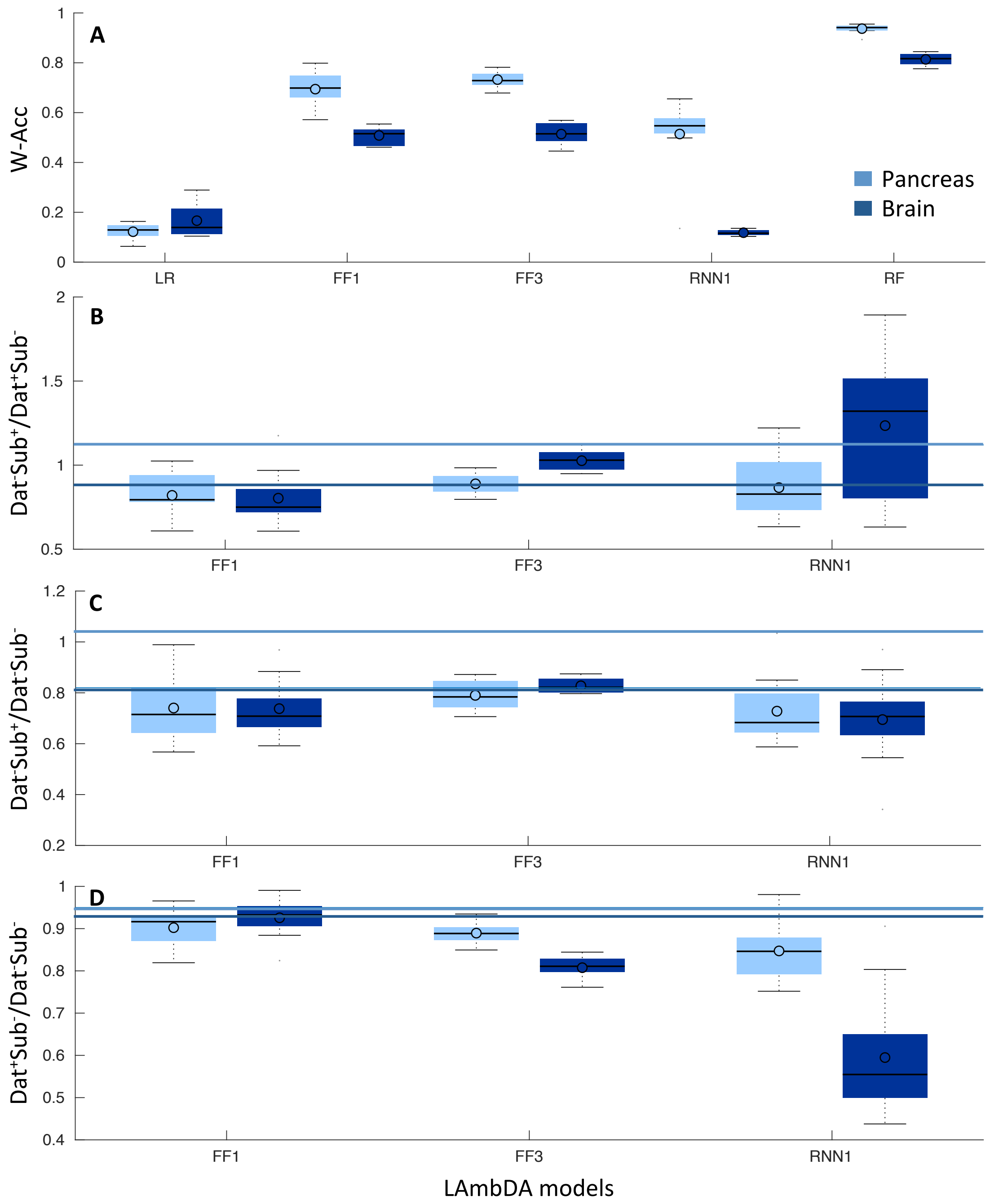


**Supplementary Fig. S3.** The boxplots of correlation and cluster distance metrics for the LAmbDA-based method and full set features. A) Unambiguous weighted accuracy with 10 rounds of cross validation. B-D) The distance ratios after feature reduction in the neural network hidden layers (LR and RF are not included because they contain no hidden layer) B) Different dataset same subtype to same dataset different subtype ratio, which indicates the level of batch effect removal for each model. C) Different dataset same subtype to different dataset different subtype, which represents the level of subtype dissimilarity. D) Same dataset different subtype to different dataset different subtype ratio, which indicate the level of noise introduced. The horizontal lines signify distance metrics using the raw data as comparison. LR: logistic regression; FF1: feedforward neural network with one layer; FF3 feedforward neural network with three layers; RNN1: recurrent neural network with one recurrent layer; and RF: random forest. W-Acc: weighted accuracy.

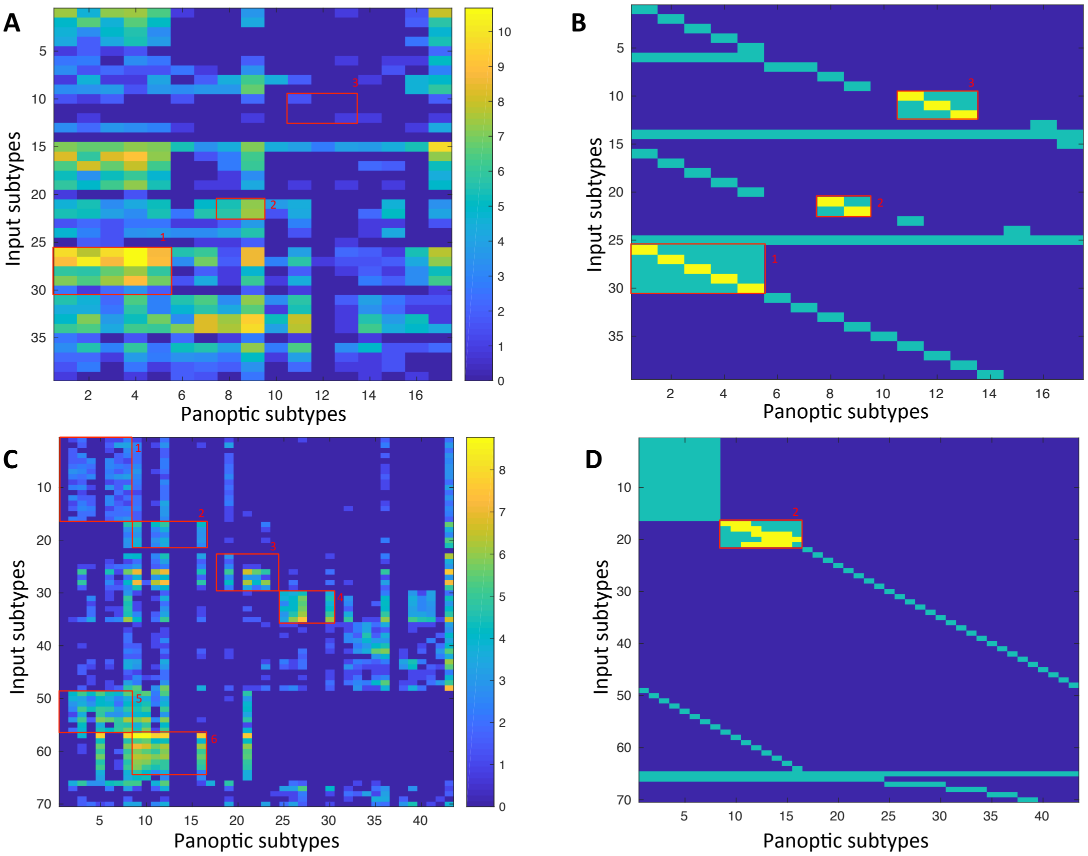


**Supplementary Fig. S4.** Confusion matrices with their associated label masks used during training for LAmbDA-LR. Each box with number indicates an AOI. A,C) Confusion matrix across three datasets where rows are original cell types and the columns are the panoptic cell types (I.e. LAmbDA output labels) for pancreas (A) and brain (C). B,D) The label mask used during LAmbDA training. Green indicates the mask used as input and yellow indicate the true labels, which were either known or inferred from the literature. C) Yellow inside of AOI1-3 indicate true labels from the starting datasets. D) Yellow indicate the cortical layer specific mapping that was inferred from each dataset's publication.

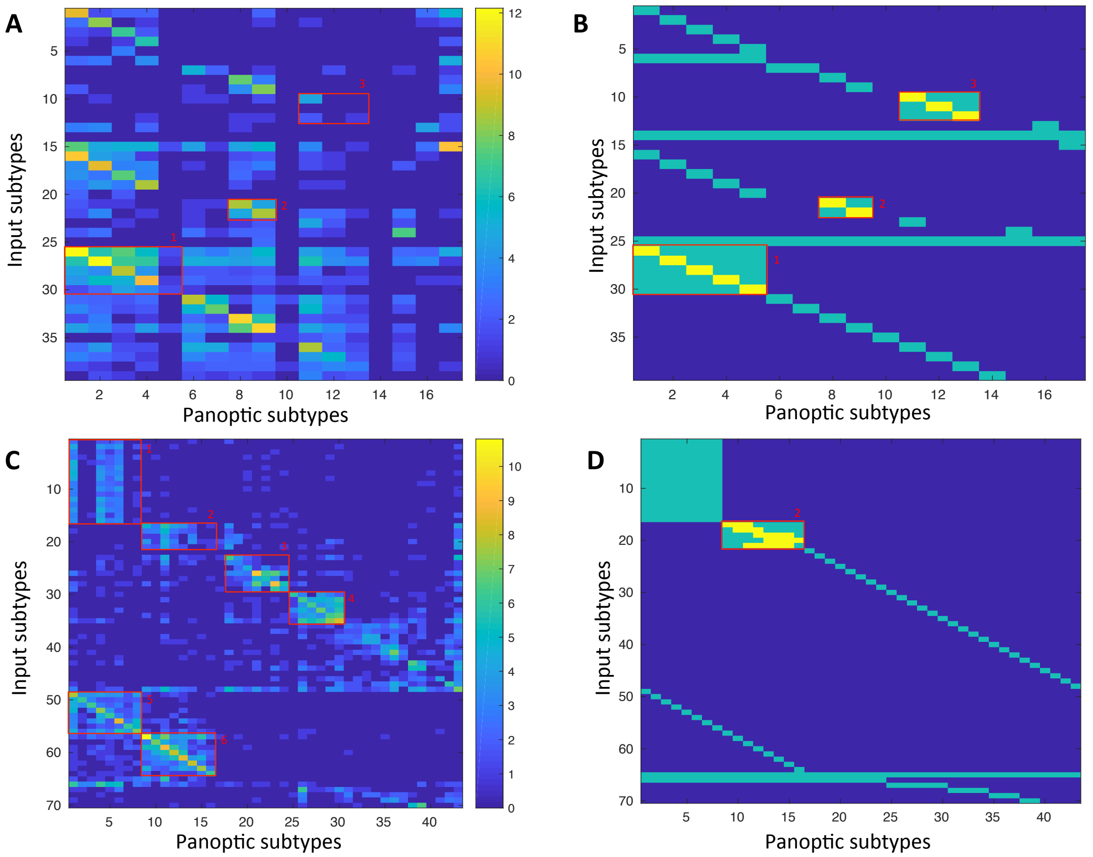


**Supplementary Fig. S5.** Confusion matrices with their associated label masks used during training for LAmbDA-FF3. Each box with number indicates an AOI. A,C) Confusion matrix across three datasets where rows are original cell types and the columns are the panoptic cell types (I.e. LAmbDA output labels) for pancreas (A) and brain (C). B,D) The label mask used during LAmbDA training. Green indicates the mask used as input and yellow indicate the true labels, which were either known or inferred from the literature. C) Yellow inside of AOI1-3 indicate true labels from the starting datasets. D) Yellow indicate the cortical layer specific mapping that was inferred from each dataset's publication.

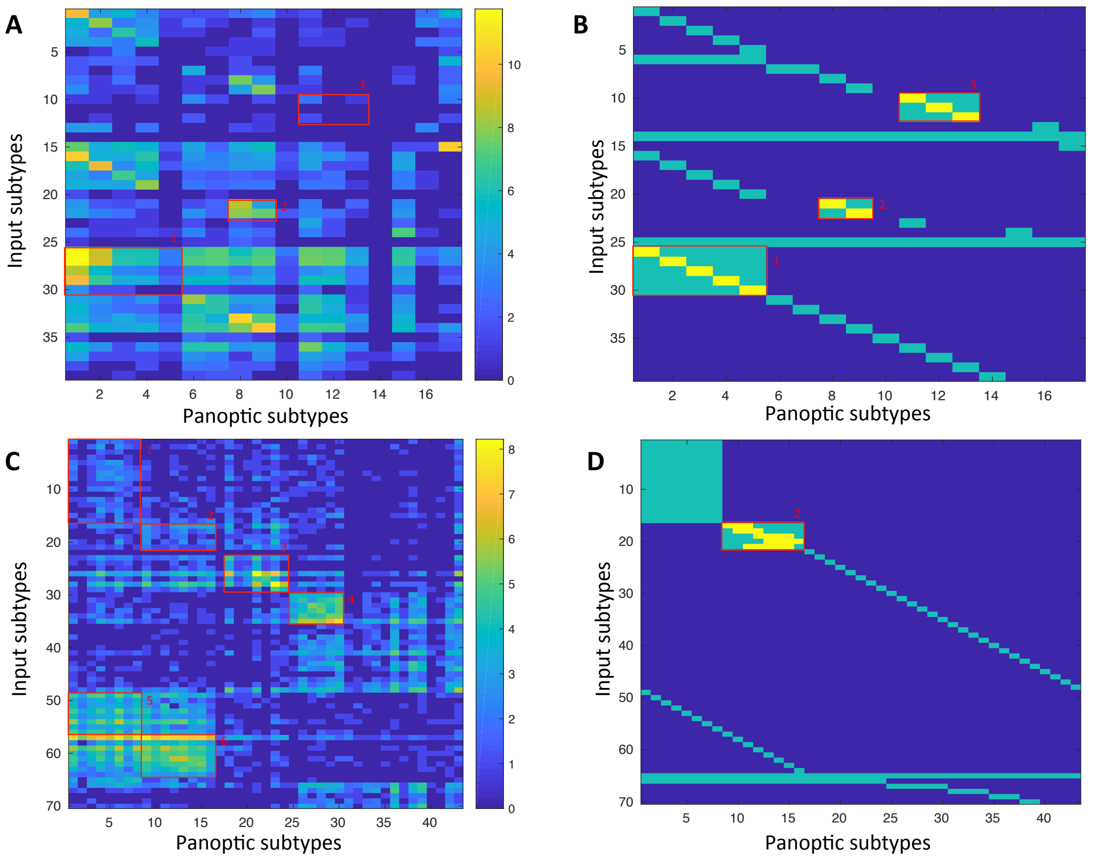


**Supplementary Fig. S6.** Confusion matrices with their associated label masks used during training for LAmbDA-RNN1. Each box with number indicates an AOI. A,C) Confusion matrix across three datasets where rows are original cell types and the columns are the panoptic cell types (I.e. LAmbDA output labels) for pancreas (A) and brain (C). B,D) The label mask used during LAmbDA training. Green indicates the mask used as input and yellow indicate the true labels, which were either known or inferred from the literature. C) Yellow inside of AOI1-3 indicate true labels from the starting datasets. D) Yellow indicate the cortical layer specific mapping that was inferred from each dataset's publication.

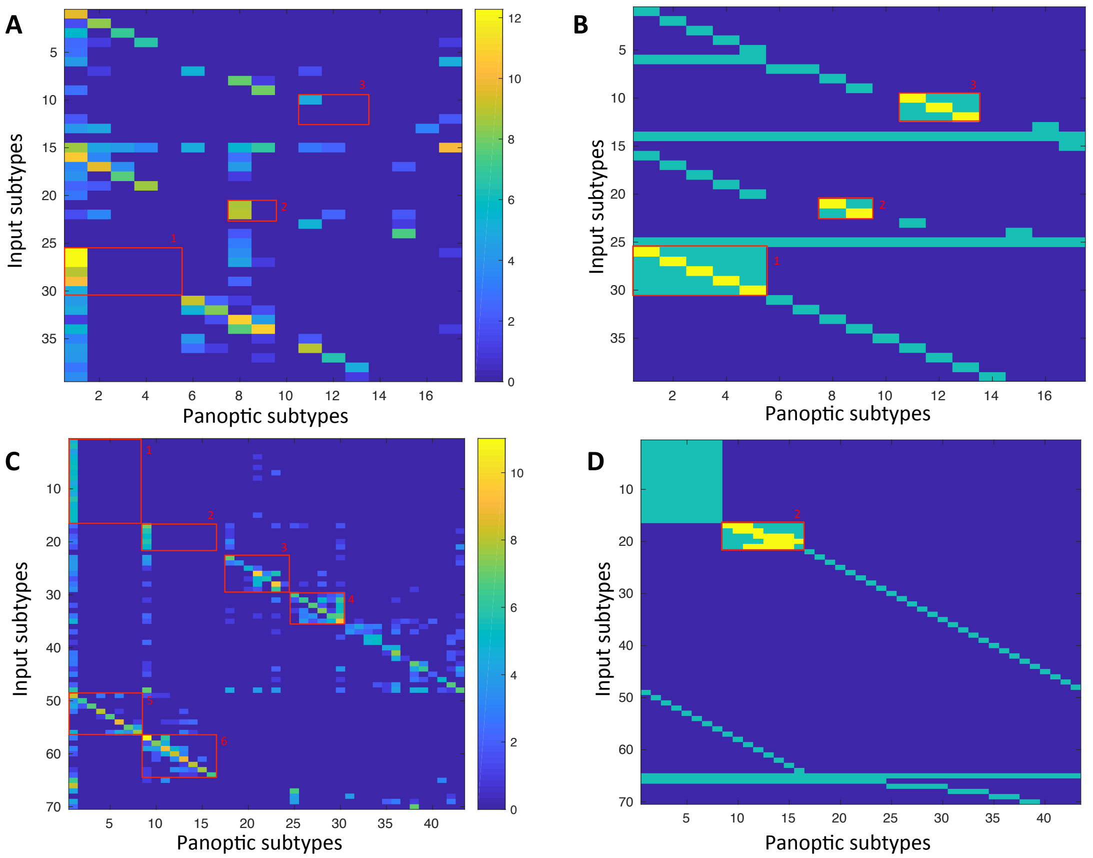


**Supplementary Fig. S7.** Confusion matrices with their associated label masks used during training for LAmbDA-RF. Each box with number indicates an AOI. A,C) Confusion matrix across three datasets where rows are original cell types and the columns are the panoptic cell types (I.e. LAmbDA output labels) for pancreas (A) and brain (C). B,D) The label mask used during LAmbDA training. Green indicates the mask used as input and yellow indicate the true labels, which were either known or inferred from the literature. C) Yellow inside of AOI1-3 indicate true labels from the starting datasets. D) Yellow indicate the cortical layer specific mapping that was inferred from each dataset's publication.

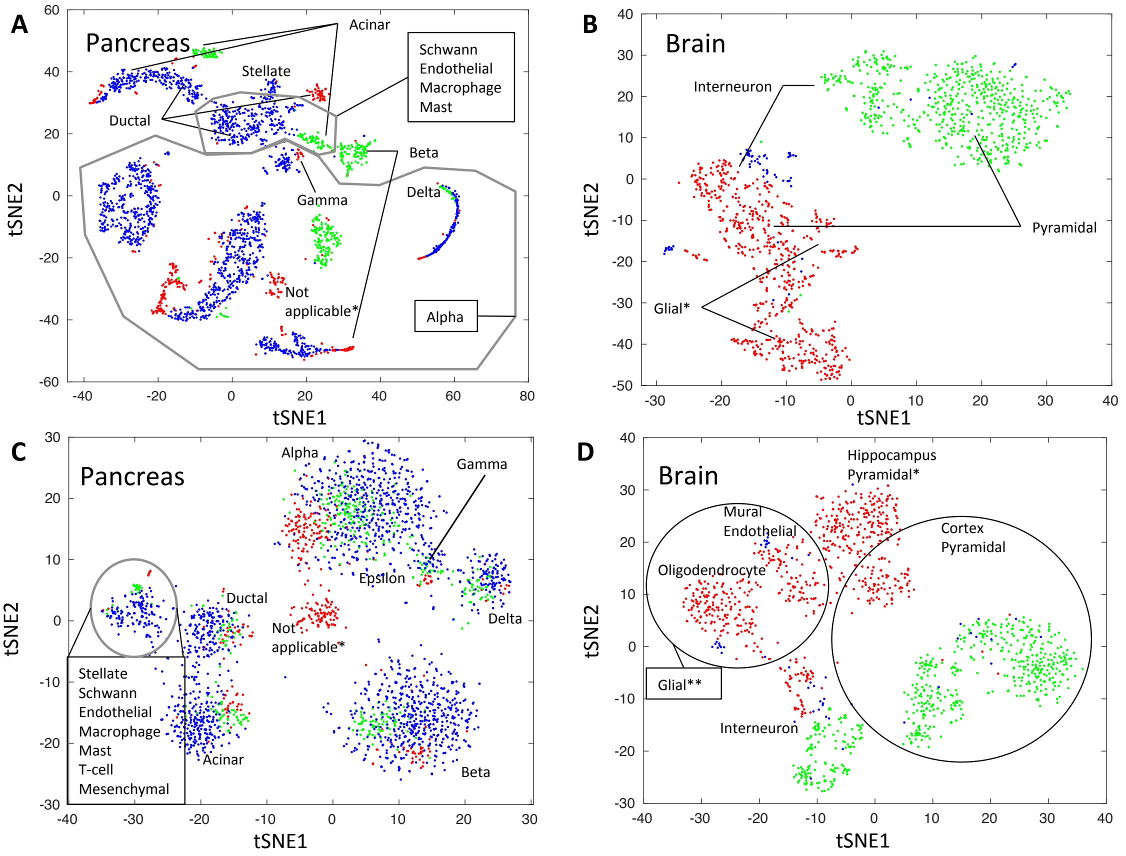


**Supplementary Fig. S8.** The tSNE dimensionality reduction from raw data using random 20% of the samples (A,B), and from the hidden layer of LAmbDA-FF3 20% of total samples contained in test set (C,D). A) Pancreatic datasets plotted using tSNE. B) Brain datasets plotted using tSNE. C) Pancreatic data from the hidden layer of LAmbDA. D) Brain data from the hidden layer of LAmbDA. The colors indicate the dataset A,C) Seger (red), Mur (green), Bar (blue), and B,D) MusNG (red), HumN (green), HumNG (blue). *Indicate cell types that are only present in one dataset. **Indicated glial cells, which are not present in the HumN dataset.

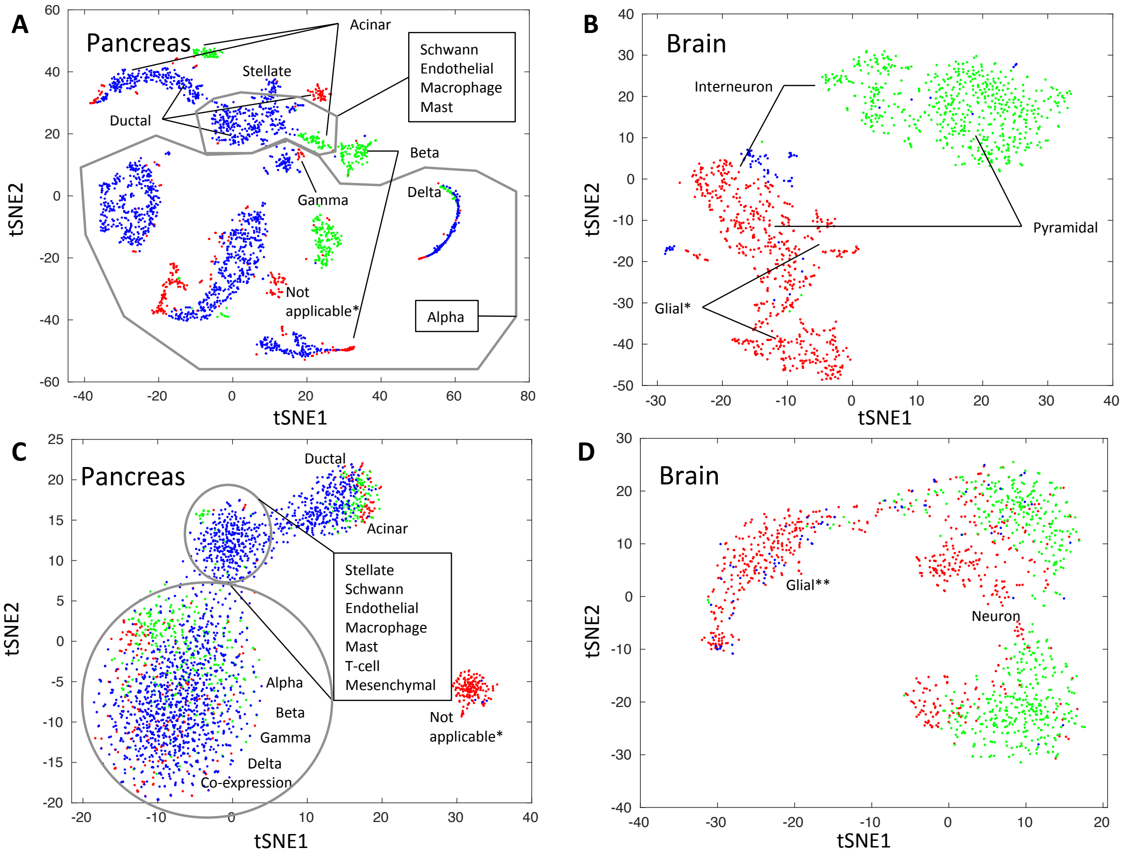


**Supplementary Fig. S9**. The tSNE dimensionality reduction from raw data using random 20% of the samples (A,B), and from the hidden layer of LAmbDA-RNN1 20% of total samples contained in test set (C,D). A) Pancreatic datasets plotted using tSNE. B) Brain datasets plotted using tSNE. C) Pancreatic data from the hidden layer of LAmbDA. D) Brain data from the hidden layer of LAmbDA. The colors indicate the dataset A,C) Seger (red), Mur (green), Bar (blue), and B,D) MusNG (red), HumN (green), HumNG (blue). *Indicate cell types that are only present in one dataset. **Indicated glial cells, which are not present in the HumN dataset.

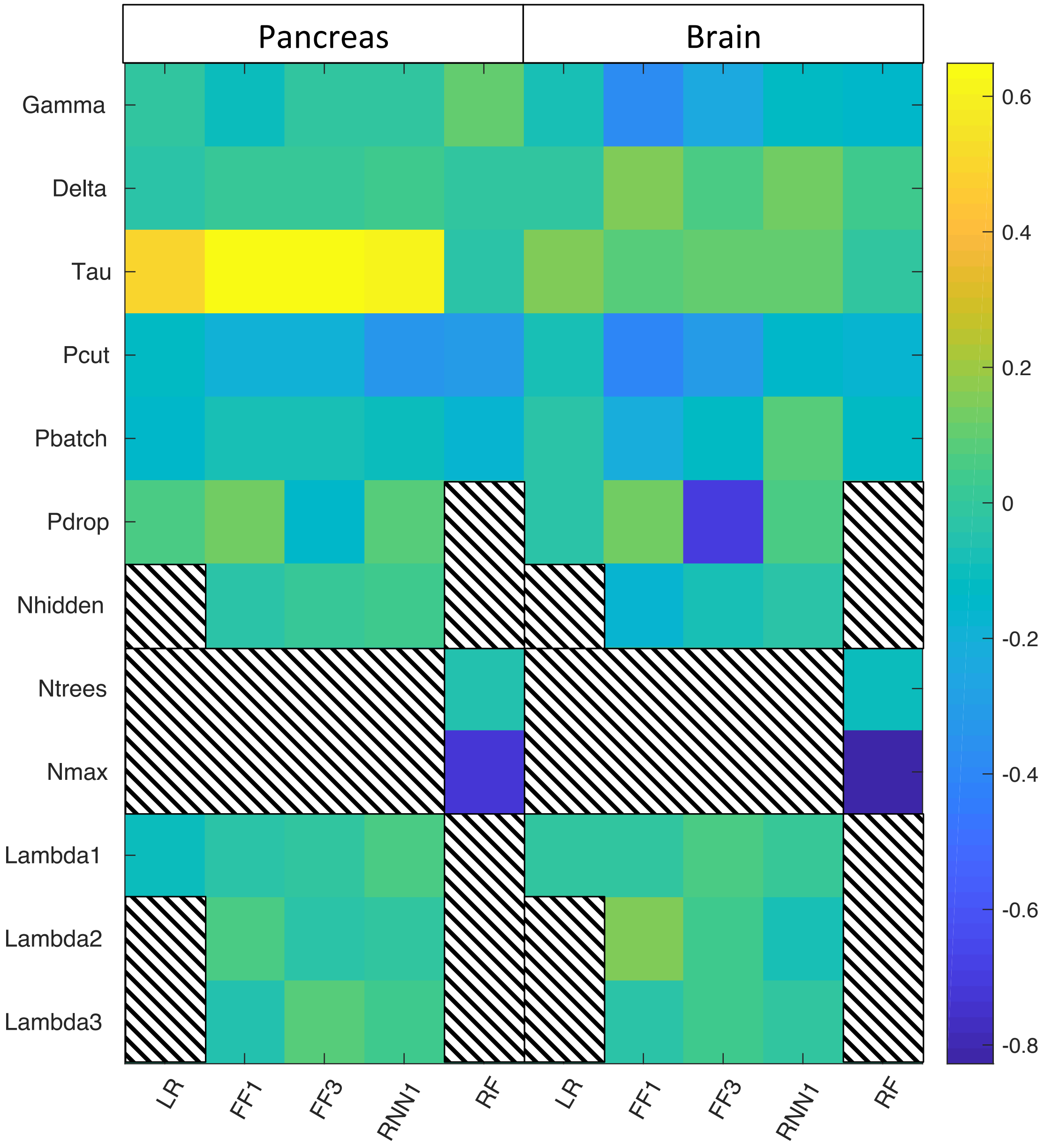


**Supplementary Fig. S10.** The correlations between hyperparameters and training error. Black and white boxes indicate hyperparameter model combinations that did not exist.

**Supplementary Table 1.** Mouse and Human Interneuron conserved subtype mapping with overlapping biomarkers from the original publications(Lake, et al., 2016; Zeisel, et al., 2015). The relationships listed in the first two columns can be seen in Figure 4C AOI1. In1-8 are the interneuron subtypes from the HumN dataset and Int1-16 are the interneuron subtypes from the MusNG dataset.

| HumN | MusNG | Biomarkers |
|---|---|---|
| In1 | Int5-8,12 | CCK,CXCL14,VIP,RELN |
| In2 | Int2,5-8,10 | CCK,CXCL14,VIP,CALB2 |
| In3 | Int6,9,10 | CCK,CXCL14,VIP,CALB2 |
| In4 | Int12-16 | CXCL14,RELN |

| | | |
|---|---|---|
| In5 | Int14-16 | |
| In6 | Int2,3,6,9 | LHX6,PVALB |
| In7 | Int1,2 | LHX6,SST,PDE1A,NPY |
| In8 | Int1,2,4,14 | LHX6,SST |

**Supplementary Table 2.** Comparison with MetaNeighbor in pancreas datasets. Cell Type column contains the cell type and dataset. The next three columns show the closest neighbors based on MetaNeighbor for each dataset. The last column contains the panoptic cell type that was identified by LAmbDA.

<table>
<tr><th></th><th colspan="3">MetaNeighbor</th><th>LAmbDA</th></tr>
<tr><th>Cell Type</th><th>Seger</th><th>Mur</th><th>Bar</th><th>Conserved</th></tr>
<tr><td>Seger|na</td><td>Seger|acinar</td><td>Mur|acinar</td><td>Bar|acinar</td><td>NA</td></tr>
<tr><td>Seger|delta</td><td>Seger|delta</td><td>Mur|delta</td><td>Bar|delta</td><td>delta</td></tr>
<tr><td>Seger|alpha</td><td>Seger|alpha</td><td>Mur|alpha</td><td>Bar|alpha</td><td>alpha</td></tr>
<tr><td>Seger|gamma</td><td>Seger|gamma</td><td>Mur|gamma</td><td>Bar|gamma</td><td>gamma</td></tr>
<tr><td>Seger|ductal</td><td>Seger|ductal</td><td>Mur|ductal</td><td>Bar|ductal</td><td>ductal</td></tr>
<tr><td>Seger|acinar</td><td>Seger|acinar</td><td>Mur|acinar</td><td>Bar|acinar</td><td>acinar</td></tr>
<tr><td>Seger|beta</td><td>Seger|beta</td><td>Mur|beta</td><td>Bar|beta</td><td>beta</td></tr>
<tr><td>Seger|endocrine</td><td>Seger|endocrine</td><td>Mur|acinar</td><td>Bar|acinar</td><td>NA</td></tr>
<tr><td>Seger|coexp</td><td>Seger|beta</td><td>Mur|beta</td><td>Bar|beta</td><td>co-expression</td></tr>
<tr><td>Seger|mhc2</td><td>Seger|unclass</td><td>Mur|mesen</td><td>Bar|macro</td><td></td></tr>
<tr><td>Seger|psc</td><td>Seger|psc</td><td>Mur|mesen</td><td>Bar|psc</td><td>stellate</td></tr>
<tr><td>Seger|endo</td><td>Seger|endo</td><td>Mur|mesen</td><td>Bar|endo</td><td>endothelial</td></tr>
<tr><td>Seger|epsilon</td><td>Seger|epsilon</td><td>Mur|epsilon</td><td>Bar|epsilon</td><td>gamma</td></tr>
<tr><td>Seger|mast</td><td>Seger|mast</td><td>Mur|mesen</td><td>Bar|mast</td><td>endothelial</td></tr>
<tr><td>Seger|unclass</td><td>Seger|unclass</td><td>Mur|mesen</td><td>Bar|psc</td><td></td></tr>
<tr><td>Mur|alpha</td><td>Seger|alpha</td><td>Mur|alpha</td><td>Bar|alpha</td><td>alpha</td></tr>
<tr><td>Mur|endo</td><td>Seger|endo</td><td>Mur|mesen</td><td>Bar|psc</td><td>endothelial</td></tr>
<tr><td>Mur|delta</td><td>Seger|delta</td><td>Mur|delta</td><td>Bar|delta</td><td>delta</td></tr>
<tr><td>Mur|beta</td><td>Seger|beta</td><td>Mur|beta</td><td>Bar|beta</td><td>beta</td></tr>
<tr><td>Mur|unclass</td><td>Seger|ductal</td><td>Mur|ductal</td><td>Bar|ductal</td><td>ductal</td></tr>
<tr><td>Mur|ductal</td><td>Seger|ductal</td><td>Mur|ductal</td><td>Bar|ductal</td><td>ductal</td></tr>
<tr><td>Mur|acinar</td><td>Seger|acinar</td><td>Mur|acinar</td><td>Bar|acinar</td><td>acinar</td></tr>
<tr><td>Mur|gamma</td><td>Seger|gamma</td><td>Mur|gamma</td><td>Bar|gamma</td><td>gamma</td></tr>
<tr><td>Mur|mesen</td><td>Seger|psc</td><td>Mur|mesen</td><td>Bar|psc</td><td>stellate</td></tr>
<tr><td>Mur|epsilon</td><td>Seger|epsilon</td><td>Mur|epsilon</td><td>Bar|epsilon</td><td>gamma</td></tr>
<tr><td>Bar|acinar</td><td>Seger|acinar</td><td>Mur|acinar</td><td>Bar|acinar</td><td>acinar</td></tr>
<tr><td>Bar|beta</td><td>Seger|beta</td><td>Mur|beta</td><td>Bar|beta</td><td>beta</td></tr>
<tr><td>Bar|delta</td><td>Seger|delta</td><td>Mur|delta</td><td>Bar|delta</td><td>delta</td></tr>
<tr><td>Bar|psc</td><td>Seger|psc</td><td>Mur|mesen</td><td>Bar|psc</td><td>stellate</td></tr>
<tr><td>Bar|ductal</td><td>Seger|ductal</td><td>Mur|ductal</td><td>Bar|ductal</td><td>ductal</td></tr>
<tr><td>Bar|alpha</td><td>Seger|alpha</td><td>Mur|alpha</td><td>Bar|alpha</td><td>alpha</td></tr>
<tr><td>Bar|epsilon</td><td>Seger|epsilon</td><td>Mur|epsilon</td><td>Bar|epsilon</td><td>gamma</td></tr>
</table>

| | | | | |
|---|---|---|---|---|
| Bar\|gamma | Seger\|gamma | Mur\|gamma | Bar\|gamma | gamma |
| Bar\|endo | Seger\|endo | Mur\|mesen | Bar\|endo | endothelial |
| Bar\|macro | Seger\|mhc2 | Mur\|mesen | Bar\|macro | mhc2 |
| Bar\|schwann | Seger\|unclass | Mur\|mesen | Bar\|schwann | stellate |
| Bar\|mast | Seger\|mast | Mur\|mesen | Bar\|mast | stellate |
| Bar\|tcell | Seger\|psc | Mur\|mesen | Bar\|tcell | endothelial |

**Supplementary Table 3.** Comparison with MetaNeighbor in brain datasets. Cell Type column contains the cell type and dataset. The next three columns show the closest neighbors based on MetaNeighbor for each dataset. The last column contains the panoptic cell type that was identified by LAmbDA.

| | MetaNeighbor | | | LAmbDA |
|---|---|---|---|---|
| Cell Type | MusNG | HumN | HumNG | Panoptic |
| MusNG\|Int10 | MusNG\|Int10 | HumN\|In2 | HumNG\|Neu | In1 |
| MusNG\|Int6 | MusNG\|Int6 | HumN\|In2 | HumNG\|Neu | In1 |
| MusNG\|Int9 | MusNG\|Int7 | HumN\|In2 | HumNG\|Neu | In1 |
| MusNG\|Int2 | MusNG\|Int2 | HumN\|In2 | HumNG\|Neu | In8 |
| MusNG\|Int4 | MusNG\|Int7 | HumN\|In2 | HumNG\|Neu | In8 |
| MusNG\|Int1 | MusNG\|Int1 | HumN\|In6 | HumNG\|Neu | In8 |
| MusNG\|Int3 | MusNG\|Int3 | HumN\|In6 | HumNG\|Neu | In6 |
| MusNG\|Int13 | MusNG\|Int13 | HumN\|In5 | HumNG\|Neu | In4 |
| MusNG\|Int16 | MusNG\|Int16 | HumN\|In4 | HumNG\|Neu | In4 |
| MusNG\|Int14 | MusNG\|Int14 | HumN\|In4 | HumNG\|Neu | In4 |
| MusNG\|Int11 | MusNG\|Int12 | HumN\|In2 | HumNG\|Neu | In1 |
| MusNG\|Int5 | MusNG\|Int5 | HumN\|In4 | HumNG\|Neu | In1 |
| MusNG\|Int7 | MusNG\|Int7 | HumN\|In2 | HumNG\|Neu | In1 |
| MusNG\|Int8 | MusNG\|Int8 | HumN\|In2 | HumNG\|Neu | In1 |
| MusNG\|Int12 | MusNG\|Int12 | HumN\|In2 | HumNG\|Neu | In1 |
| MusNG\|Int15 | MusNG\|Int16 | HumN\|In4 | HumNG\|Neu | In4 |
| MusNG\|none | MusNG\|Astro1 | HumN\|Ex3 | HumNG\|Endo | none |
| MusNG\|S1PyrL4 | MusNG\|S1PyrL4 | HumN\|Ex3 | HumNG\|Neu | Ex3 |
| MusNG\|ClauPyr | MusNG\|ClauPyr | HumN\|Ex8 | HumNG\|Neu | |
| MusNG\|S1PyrL5 | MusNG\|S1PyrL5 | HumN\|Ex5 | HumNG\|Neu | Ex5 |
| MusNG\|S1PyrL23 | MusNG\|S1PyrL23 | HumN\|Ex1 | HumNG\|Neu | Ex3 |
| MusNG\|S1PyrDL | MusNG\|S1PyrDL | HumN\|Ex5 | HumNG\|Neu | S1PyrDL |
| MusNG\|S1PyrL5a | MusNG\|S1PyrL5a | HumN\|Ex3 | HumNG\|Neu | Ex3 |
| MusNG\|SubPyr | MusNG\|SubPyr | HumN\|Ex5 | HumNG\|Neu | SubPyr |
| MusNG\|CA1Pyr1 | MusNG\|CA1Pyr1 | HumN\|Ex1 | HumNG\|Neu | CA1Pyr1 |
| MusNG\|S1PyrL6b | MusNG\|S1PyrL6b | HumN\|Ex5 | HumNG\|Neu | S1PyrDL |
| MusNG\|S1PyrL6 | MusNG\|S1PyrL6 | HumN\|Ex5 | HumNG\|Neu | Ex6 |
| MusNG\|CA1Pyr2 | MusNG\|CA1Pyr2 | HumN\|Ex1 | HumNG\|Neu | CA1Pyr2 |
| MusNG\|CA1PyrInt | MusNG\|CA1Pyr1 | HumN\|In3 | HumNG\|Neu | CA1Pyr1 |

| | | | | |
|---|---|---|---|---|
| MusNG\|CA2Pyr2 | MusNG\|CA2Pyr2 | HumN\|Ex5 | HumNG\|Neu | CA2Pyr2 |
| MusNG\|Oligo1 | MusNG\|Oligo1 | HumN\|Ex8 | HumNG\|Oligo | Oligo1 |
| MusNG\|Oligo3 | MusNG\|Oligo3 | HumN\|Ex2 | HumNG\|Oligo | Oligo3 |
| MusNG\|Oligo4 | MusNG\|Oligo3 | HumN\|Ex2 | HumNG\|Oligo | Oligo4 |
| MusNG\|Oligo2 | MusNG\|Oligo3 | HumN\|Ex2 | HumNG\|Oligo | Oligo2 |
| MusNG\|Oligo6 | MusNG\|Oligo3 | HumN\|Ex2 | HumNG\|Oligo | Oligo6 |
| MusNG\|Oligo5 | MusNG\|Oligo3 | HumN\|Ex2 | HumNG\|Oligo | Oligo6 |
| MusNG\|Mgl1 | MusNG\|Mgl1 | HumN\|In2 | HumNG\|Micro | Pvm1 |
| MusNG\|Mgl2 | MusNG\|Pvm1 | HumN\|In2 | HumNG\|Micro | Pvm1 |
| MusNG\|Pvm1 | MusNG\|Pvm1 | HumN\|In8 | HumNG\|Micro | Pvm1 |
| MusNG\|Pvm2 | MusNG\|Pvm2 | HumN\|In2 | HumNG\|Micro | Pvm1 |
| MusNG\|Vsmc | MusNG\|Vsmc | HumN\|Ex8 | HumNG\|Endo | |
| MusNG\|Vend2 | MusNG\|Vend2 | HumN\|Ex1 | HumNG\|Endo | Vend2 |
| MusNG\|Peric | MusNG\|Vend2 | HumN\|Ex1 | HumNG\|Endo | Vend2 |
| MusNG\|Vend1 | MusNG\|Vend1 | HumN\|Ex8 | HumNG\|Endo | Vend2 |
| MusNG\|Astro2 | MusNG\|Astro1 | HumN\|Ex2 | HumNG\|Astro | Astro2 |
| MusNG\|Astro1 | MusNG\|Astro1 | HumN\|Ex1 | HumNG\|Astro | Astro1 |
| MusNG\|Choriod | MusNG\|Astro1 | HumN\|Ex2 | HumNG\|Astro | none |
| MusNG\|Epend | MusNG\|Epend | HumN\|Ex1 | HumNG\|Endo | Epend |
| HumN\|Ex1 | MusNG\|CA1Pyr1 | HumN\|Ex1 | HumNG\|non | Ex1 |
| HumN\|Ex3 | MusNG\|S1PyrL4 | HumN\|Ex3 | HumNG\|non | Ex3 |
| HumN\|Ex4 | MusNG\|S1PyrL4 | HumN\|Ex4 | HumNG\|non | Ex4 |
| HumN\|In6 | MusNG\|Int13 | HumN\|In6 | HumNG\|Neu | In6 |
| HumN\|In1 | MusNG\|Int12 | HumN\|In1 | HumNG\|Neu | In1 |
| HumN\|Ex5 | MusNG\|S1PyrL5 | HumN\|Ex5 | HumNG\|non | Ex5 |
| HumN\|In5 | MusNG\|Int14 | HumN\|In5 | HumNG\|Neu | In5 |
| HumN\|Ex7 | MusNG\|S1PyrL5 | HumN\|Ex7 | HumNG\|non | Ex7 |
| HumN\|In8 | MusNG\|Int5 | HumN\|In8 | HumNG\|non | In8 |
| HumN\|Ex2 | MusNG\|S1PyrL4 | HumN\|Ex2 | HumNG\|non | Ex2 |
| HumN\|Ex6 | MusNG\|SubPyr | HumN\|Ex6 | HumNG\|non | Ex6 |
| HumN\|In7 | MusNG\|Int2 | HumN\|In7 | HumNG\|non | In7 |
| HumN\|In4 | MusNG\|Int16 | HumN\|In4 | HumNG\|Neu | In4 |
| HumN\|Ex8 | MusNG\|ClauPyr | HumN\|Ex8 | HumNG\|non | Ex8 |
| HumN\|In3 | MusNG\|Int10 | HumN\|In3 | HumNG\|Neu | In3 |
| HumN\|In2 | MusNG\|Int10 | HumN\|In2 | HumNG\|Neu | In2 |
| HumNG\|non | MusNG\|Astro2 | HumN\|Ex8 | HumNG\|non | Ex3 |
| HumNG\|Oligo | MusNG\|Oligo3 | HumN\|Ex8 | HumNG\|Oligo | Oligo2 |
| HumNG\|Astro | MusNG\|Astro1 | HumN\|Ex2 | HumNG\|Astro | Astro2 |
| HumNG\|Micro | MusNG\|Pvm1 | HumN\|Ex8 | HumNG\|Micro | Pvm1 |
| HumNG\|Neu | MusNG\|Int8 | HumN\|In2 | HumNG\|Neu | In1 |
| HumNG\|Endo | MusNG\|Vend2 | HumN\|Ex8 | HumNG\|Endo | Vend2 |
